# LEM-3/ANKLE1 nuclease prevents the formation of syncytium between postmitotic sister cells and safeguards neuronal differentiation

**DOI:** 10.1101/2025.11.13.688214

**Authors:** Siyu Deng, Chaogu Zheng

## Abstract

In animal development, cells exit the cell cycle after a final division to differentiate into specialized types. However, the impact of failed cytokinesis on the fate specification of postmitotic cells, especially neurons, remain poorly understood. We analysed the *Caenorhabditis elegans* mutants lacking the midbody-tethered endonuclease LEM-3/ANKLE1, which resolves chromatin bridges during cytokinesis. Using genetic analysis, fluorescence microscopy, and RNA visualization, we found that unresolved DNA bridges lead to the formation of stable intercellular canals, creating binucleate syncytia between touch receptor neurons and sisters, which allows thorough cytoplasmic mixing despite independent nuclear transcription. In some cases, the cytoplasmic linkage between sister cells also led to the suppression of neuronal fates and prevention of apoptotic fates. Thus, LEM-3 prevents syncytium formation to ensure proper cellular differentiation.

## Introduction

In development, cells exit the cell cycle after the final round of cell division in order to differentiate into specific cell types. During this process, sister cells are separated by cytokinesis, so that they can adopt different cell fates. However, surprisingly little is known about the consequence of failed cytokinesis in terminal differentiation. Can cells, especially neurons, still differentiate if they are physically connected to their sister cells? How does the mixing of cytoplasmic content affect the fate specification of the two sister cells that are supposed to differentiate into distinct cell fates? In this study, we address these questions by studying the *C. elegans lem-3*(-) mutants, which caused the formation of a persistent cytoplasmic linkage (referred to as “intercellular canal”) between sister cells due to failure in resolving chromatin bridges.

LEM-3 is an evolutionarily conserved LEM domain protein that also contains N-terminal Ankyrin repeats and a C-terminal endonuclease domain. *lem-3* was initially discovered in a genetic screen for DNA repair genes, as the *lem-3*(-) mutants were hypersensitive to excessive DNA damage (*1*). More recently, LEM-3 was found to localize to the midbody in the late stages of cytokinesis to resolve chromatin bridges (or other DNA bridges) caused by incomplete DNA replication, persistent recombination intermediates, or defects in chromosome condensation or decatenation (*2*). The LEM-3 homolog in humans, ANKLE1, shows nuclease activities towards a range of branched DNA species (*3, 4*) and is similarly recruited to the midbody to process chromatin bridges (*5*), suggesting conserved functions across species. The formation of chromatin bridges delays cytokinesis (*6, 7*), but persistent bridges are eventually cleaved by actomyosin contractile force in continuously dividing cells (*8*). However, it remains unclear whether the chromatin bridges would be broken and the intercellular canal be severed following the terminal cell division that generates two postmitotic daughter cells.

In certain tissues, multinucleate syncytium forms normally in development. For example, the *C. elegans* hypodermis and seam cells are multinucleate syncytia that are generated by cell fusion; in particular, the hyp7 cell is a syncytium containing 139 nuclei (*9*). In another example, the germline of *C. elegans* hermaphrodites are made of syncytial structures that contain more than 1000 nuclei as a result of incomplete cytokinesis (*10*). Muscles also form multinucleated syncytia in various organisms. In the above cases, syncytia connect the same types of cells. The developmental fate of syncytia that connect different types of cells is less clear. Here, we studied this question using the binucleate syncytia formed between sister cells due to unresolved chromatin bridges in *lem-3*(-) mutants.

## Results

### The loss of *lem-3* results in binucleate syncytia connecting TRNs and sister cells through an intercellular canal

In a forward genetic screen searching for genes that regulate TRN fate specification, we isolated a mutant allele *unk14* that caused the expression of TRN markers in extra cells. The wild-type animals have six TRNs, including ALML/R, AVM, PVM, and PLML/R, which were specifically labelled by the *mec-17p::TagRFP* transgene, whereas *unk14* mutants showed extra cells near AVM and PLM cell bodies in ∼15% of the animals at 25°C. Strikingly, the extra cells were physically connected with the AVM and PLM cell body through an intercellular canal, which is also marked by TagRFP (Fig. 1A-D). Using chromosomal mapping and whole genome resequencing, we mapped *unk14* to the V618M mutation in *lem-3*, which codes for a midbody-tethered, LEM domain-containing DNA endonuclease (*2, 11*). To confirm that the loss of *lem-3* caused the phenotype of producing extra TRN-like cells, we tested a nonsense allele of *lem-3*, named *mn155(R190*)*, which caused similar phenotype and failed to complement *unk14*. We also generated *lem-3* knockout alleles (*unk84* and *unk85*) using CRISPR/Cas9-mediated gene editing and found that these alleles also led to the presence of extra cells labelled by TagRFP. Importantly, the extra TRN phenotype in *lem-3*(-) mutants was confirmed using other TRN fate markers (Fig. S1A-B) and can be rescued by transgenes carrying a wild-type copy of *lem-3(+)*, further confirming that the loss of *lem-3* resulted in developmental defects in restricting TRN fate in the six canonical TRNs (Fig. 1E).

**Figure 1.**
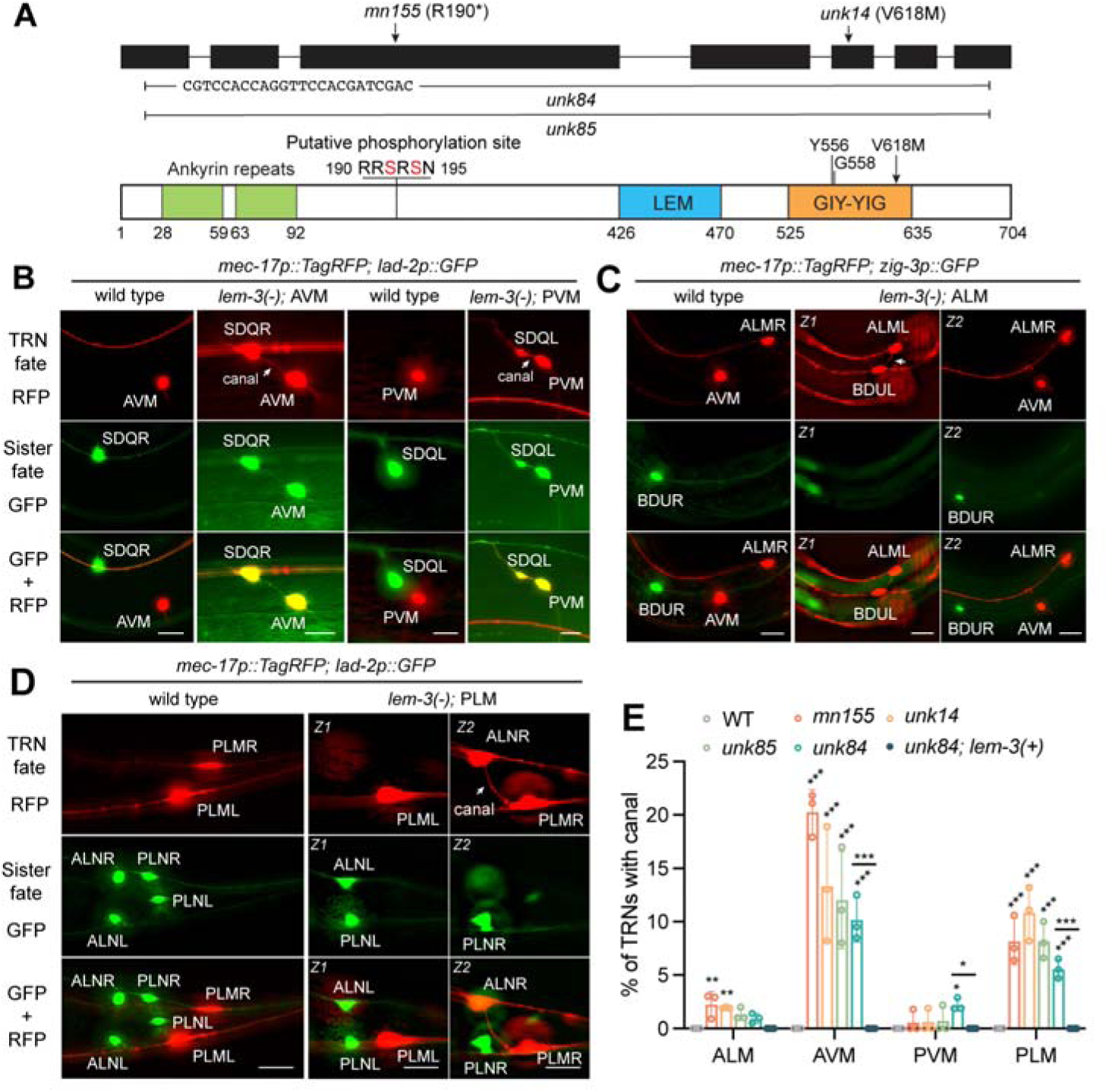
Mutations in *lem-3* lead to the formation of syncytia between TRNs and their sister cells. (A) Schematic cartoon of the *lem-3* gene structure and LEM-3 protein domain structure, as well as the molecular lesion of the *lem-3* alleles used. (B) The presence of both TRN marker *uIs115[mec-17p::TagRFP]* and SDQL/R marker *uIs130[lad-2p::GFP]* in the AVM-SDQR and PVM-SDQL syncytia in *lem-3(unk84)* mutants. Arrows point to the intercellular canal connecting the two cell bodies. Scale bars = 20 μm. (C) ALML-BDUL syncytium expresses TRN marker but does not express BDU marker *otIs14[zig-3p::GFP]* in *lem-3(unk84)* mutants. “Z1” and “Z2” mean two focal planes of the same animal. (D) PLMR-ALNR syncytium expresses TRN marker but not the ALN marker *uIs130[lad-2p::GFP]* (which also labels PLN). (E) The penetrance of binucleate syncytium connected by the canal for the four TRN subtypes at 25°C for the four *lem-3* mutants. The syncytium disappeared when the wild-type *lem-3(+)* was expressed in *lem-3(unk84)* mutants. Mean ± SD is shown. Single, double, and triple asterisks indicate *p* < 0.05, 0.01, and 0.001, respectively, in a post-ANOVA Tukey’s HSD test comparing the mutants with the wild-type or selected pairs.

Previous studies found that during the early cell divisions of *C. elegans* embryos, LEM-3 is recruited to the midbody from anaphase to the late stage of cytokinesis to resolve chromatin bridges caused by various errors in DNA replication, recombination, or chromosome condensation (*2*). The presence of chromatin bridges (and possibly ultra-fine DNA bridges) delays abscission through the activation of Aurora B kinase-mediated NoCut pathway (*6*). Aurora B activates LEM-3, which cleaves the DNA bridge and allows cytokinesis to proceed. In *lem-3*(-) mutants, chromatin bridges are not processed and are eventually broken by actomyosin dependent-mechanical force, which results in error-prone repair and chromosomal rearrangement (*12*). This forced break of chromatin bridges by cytokinesis were found in continuously dividing cells in culture or in the early embryos. However, the persistence of the intercellular canals connecting TRNs and presumably their sister cells in adult animals (>3 days after the scheduled cell division) led us to hypothesize that cytokinesis was not only delayed but aborted in response to the chromatin bridges in the last round of cell division that generates postmitotic cells.

We confirmed that the extra cells labelled by the TRN markers are the sister cells of TRNs by examining the cellular features for each of the four TRN subtypes in the *lem-3*(-) mutants. For the postembryonic AVM neuron, the extra cell connected with it resembled the morphology of its sister cell SDQR neuron (Fig. S1C), contained a nucleus labelled by histones (Fig. S1D), and expressed SDQ markers *lad-2p::GFP*, *flp-12p::GFP*, *ast-1::GFP*, and *ser-2p2::GFP* (Fig. 1B and S1E). Interestingly, AVM was also labelled by the SDQ marker, suggesting the co-expression of cell fate markers in the binucleate syncytium. Similar results were obtained for PVM-SDQL syncytium, although its occurrence has much lower penetrance (<3%) than the AVM-SDQR syncytium (10-20%) (Fig. 1B and S1F).

The sister cell of ALM is the peptidergic interneuron BDU. We observed the ALM-BDU syncytium in ∼5% of the *lem-3*(-) mutants. In some cases, the ALM cell body is directly connected with the BDU cell body (Fig. 1C), while in other cases, a canal linked the ALM cell body or anterior neurite with the BDU posterior neurite (Fig. S1A or S2A-C). Despite that ALM and BDU migrate towards the opposite directions after the cell division, their physical connection was maintained throughout development into adulthood and reached >100 μm in length in adults. BDU markers *zig-3p::GFP*, *nlp-80p::GFP*, *C15C7.4p::GFP*, and *flp-10p::GFP* were not expressed in the ALM-BDU syncytium (Fig. 1C and S2A-B), suggesting that the BDU fate was suppressed by the linkage with ALM. Nevertheless, we found exceptions in two BDU markers *ceh-14::EGFP* and *ser-2p2::GFP*, whose expression persisted in the ALM-BDU syncytium, suggesting that the suppression of BDU fate may be incomplete (Fig. 1B and S2A-B). In fact, the morphology of the ALM-connected BDU largely resembled the normal BDU, indicating at least partial differentiation of the BDU fate (Fig. S2C).

In the case of PLM, the TRN fate also dominated over the fate of its sister cell, ALN neuron, when the two cells were connected. The PLM-ALN syncytium was only labelled by the TRN markers (e.g., *mec-17p::TagRFP*, *mec-4p::YFP*, and *mec-18p::GFP*) but not by the ALN markers (e.g., *lad-2p::GFP*, *nlp-20p::GFP*, *ast-1::GFP*, and *ser-2p2::GFP*) in *lem-3*(-) mutants (Fig. 1C, S1A, and S2D-E). Whenever an extra cell was labelled by the TRN marker, we observed the loss of a GFP-labelled ALN neuron, which indicated that the formation of PLM-ALN connection converted the ALN neuron into PLM-like cells.

### Ectopic presence of MEC-3 converts TRN sisters into TRN-like fate

The suppression of BDU and ALN fates in the ALM-BDU and PLM-ALN syncytia, respectively, could be explained by the presence of the TRN fate selector MEC-3 (*13*) in the sister neurons *via* the diffusion through the intercellular canal (see Figure 3). To test this idea, we expressed *mec-3* in all neurons using the *unc-119* promoter and found that the forced expression of *mec-3* in BDU and ALN neurons indeed activated the TRN fate marker and suppressed the sister fate markers (Fig. S3A-B), confirming that ectopic presence of MEC-3 in the sister cells can convert them into a TRN-like state. Interestingly, the converted BDU neuron had an ALM-like morphology (e.g., posteriorly displaced cell body, subdorsal axons, and the loss of ventral dip) and did not express any BDU markers (e.g., *zig-3p::GFP* and *ser-2p2::GFP*; Fig. S3C), suggesting that the fate conversion caused by MEC-3 overexpression is more extensive than that induced by MEC-3 diffusion from the ALM cell body in the ALM-BDU syncytium. Our results are consistent with previous finding that MEC-3 could outcompete the BDU fate selector PAG-3 for binding to the master regulator UNC-86, which is required for the specification of both ALM and BDU fates (*14*). Although the ALN fate selector is unknown, we reason that a similar mechanism might allow ectopic MEC-3 to transform ALN into a TRN-like cell.

### Independent transcriptional programs of the two nuclei in the syncytium

The co-labelling of AVM-SDQR and PVM-SDQL syncytia by both TRN and the sister cell fate markers could result from either 1) active transcription of both fate reporters in the two nuclei of the syncytia due to the mixing of cell fate determinants or 2) the diffusion of fluorescent proteins produced only in one cell body. To discern the two possibilities, we visualize the transcription of the cell fate-associated genes from their endogenous loci in the binucleate syncytia using single-molecule fluorescence in situ hybridization (smFISH). In the majority of the AVM-SDQR syncytium, only the AVM cell body contained the mRNAs of TRN genes (e.g., *mec-17*, *mec-18*, and *mec-3*), while only the SDQR cell body contained the mRNAs of SDQ genes (*lad-2*, *ast-1*, *flp-12*, and *ser-2*; Fig. 2A, S4, and S5), suggesting that the two cells transcribed different genetic programs despite the cytoplasmic connection. This transcriptional independence was not due to the lack of selector diffusion (see below) but the inability of the selectors to activate their downstream genes in the nucleus of the sister cell. We reason that restrictive chromatin state, as previously suggested (*15, 16*), may have prevented the activation of alternative fate programs by the ectopic terminal selectors and avoided the mixing of cell fate at the transcriptional level in most AVM-SDQR syncytia.

**Figure 2.**
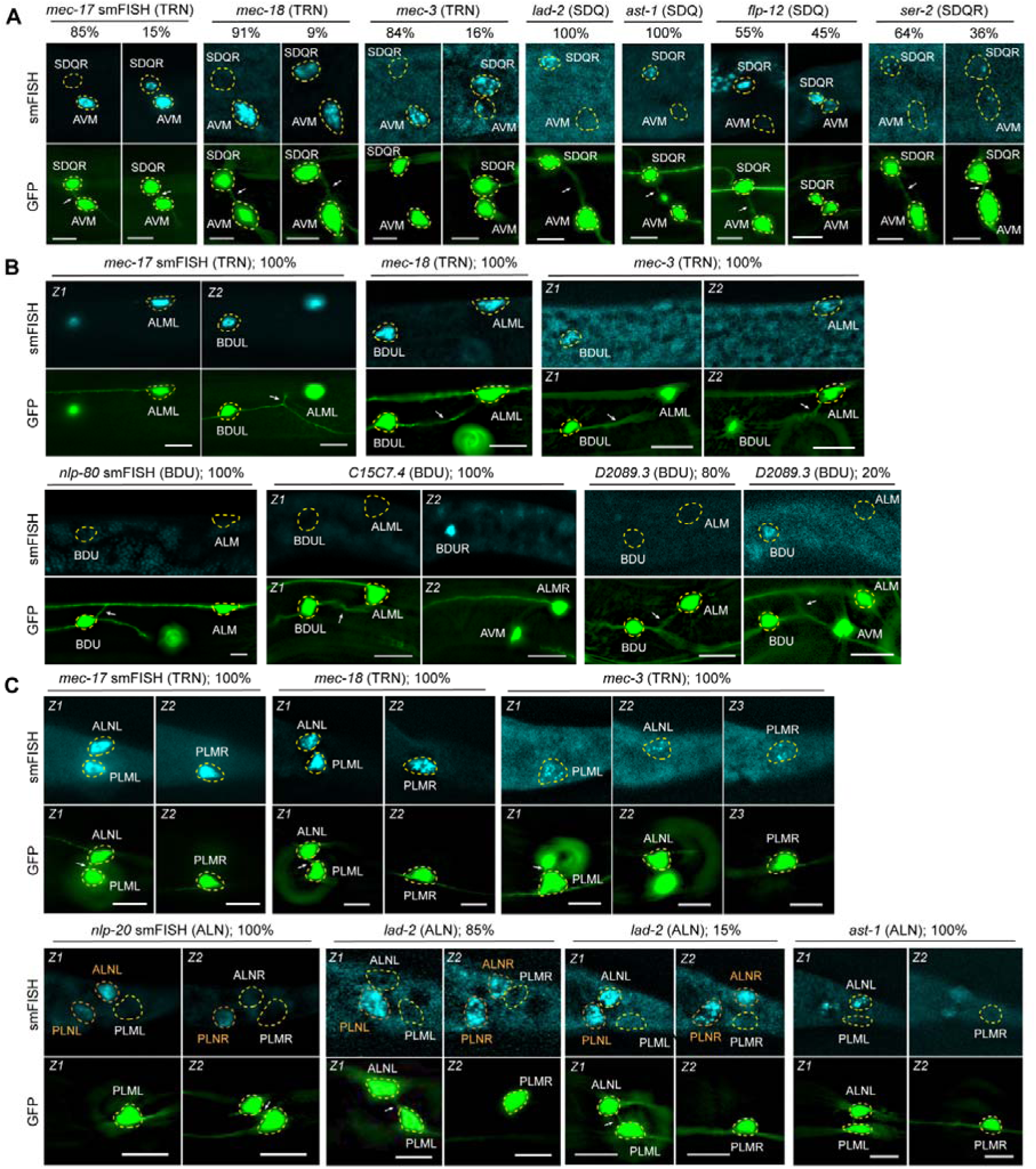
Two connected cell bodies in the syncytium have distinct transcriptional programs. (A) smFISH signals for the endogenous mRNAs of TRN fate markers (*mec-17*, *mec-18*, and *mec-3*) and SDQ markers (*lad-2*, *ast-1*, *flp-12*, and *ser-2*) in the AVM-SDQR syncytia. *ser-2* was used as SDQR but not SDQL marker, since its RNAs were found in WT PVM (the sister of SDQL). *lem-3(unk84)* mutants were used; *uIs31[mec-17p::GFP]* was used to label the TRNs and the syncytia formed between TRNs and their sisters. The percentage of the two different phenotypes were shown; at least 20 syncytia were imaged for each probe. Dashed circles indicate the outline of the cell; arrows point to the intercellular canal. Scale bars = 10 μm. The same applies to other panels. (B) smFISH signals for the mRNAs of TRN and BDU markers (*nlp-80*, *C15C7.4*, and *D2089.3*) in the two cell bodies of the ALM-BDU syncytia. (C) smFISH signals for TRN and ALN markers (*nlp-20*, *lad-2,* and *ast-1*) in the two cell bodies of the PLM-ALN syncytia. “Z1” and “Z2” mean two focal planes of the same animal.

However, in about 15% of the animals, we did observe both nuclei of the syncytium actively transcribing the TRN genes, suggesting that the TRN and SDQ fates could be co-expressed from the same nucleus in some cases (Fig. 2A). This result is consistent with the finding that forced overexpression of *mec-3* in SDQR can turn on the TRN fate without repressing the SDQR fate (Fig. S3A and S3C), suggesting that high level of ectopic MEC-3 can enable the co-existence of TRN and SDQ fate in the same cell. Interestingly, two SDQ fate markers *flp-12* and *ser-2* were activated in ∼40% of the AVM nuclei, suggesting that some AVM may adopt a partially mixed fate in the syncytium. The PVM-SDQL syncytium was similar to the AVM-SDQR pair but with a higher frequency of cell fate mixing (Fig. S6A).

For the ALM-BDU syncytium, we found that both nuclei transcribed endogenous TRN fate markers in all cases, and the expression of BDU fate markers (e.g., *nlp-80*, *C15C7.4*, *D2089.3*, *ser-2*) were suppressed to various degrees (Fig. 2B, S4-S6), which supports the conversion of BDU into a TRN-like fate due to the diffusion of MEC-3. A similar fate conversion of ALN neuron was observed in the PLM-ALN syncytium, since both cell bodies showed the mRNAs for TRN genes and did not contain the mRNAs for the ALN genes (e.g., *nlp-20*, *lad-2*, and *ser-2*) (Fig. 2C), which confirmed the suppression of ALN fate in the syncytium due to the diffusion of MEC-3 from PLM to the ALN. Interestingly, the ALN gene *ast-1* persisted in the ALN nucleus of all the PLM-ALN syncytia, suggesting that the fate conversion is still partial (Fig. 2C). It is worth noting that although the *otIs358[ser-2p2::GFP]* reporter is not expressed in wild-type ALM and PLM neurons, we detected *ser-2* mRNAs in ∼60% of the TRNs, making it a suboptimal marker for the sister fate (Fig. S4C and S4D). Nevertheless, we still observed the suppression of *ser-2* mRNA expression in 13% and 38% of ALM-BDU and PLM-ALN syncytia, respectively, compared to their 100% presence in the wild-type BDU and ALN neurons (Fig. S6B and S6C). The above results suggest that the transcriptomes of the two nuclei in a syncytium are not necessarily synchronized and can be independently regulated. The fate specification programs of the TRNs and their sister cells appear to have different susceptibilities to perturbation caused by the cytoplasmic linkage between them.

### The mixing of cellular contents between two physically connected cells

The AVM-SDQR syncytium provided an opportunity to understand the extent of cytoplasmic mixing between two transcriptionally (and presumably also translationally) independent but physically connected cells. When labelled by the *mec-17p::GFP* reporter, GFP mRNA was only detected in the AVM cell body of the syncytia (Fig. 3A and S6D), but the GFP protein diffused to fill the entire syncytium, including neuronal processes, suggesting cytoplasmic soluble proteins can pass through the intercellular canal. To confirm this idea, we expressed the photoconvertible Dendra2 from the *mec-17* promoter in the AVM-SDQR syncytium, photoconverted Dendra2 from green to red only in the AVM cell body, and observed the red fluorescent signal in the SDQR cell bodies within minutes (Fig. 3B-C). Similar results were obtained in the ALM-BDU syncytium, although there is a slower kinetics in the increase of Dendra2 signals possibly due to the longer passage way connecting the ALM and BDU cell bodies (Fig. S7). Moreover, when we severed the AVM-SDQR canal using a UV laser, the TagRFP expressed from the TRN-specific *mec-17* promoter gradually disappeared in the SDQR over time, supporting that the SDQR nucleus did not activate the *mec-17* promoter and the fluorescent protein was produced in AVM and diffused into SDQR (Fig. 3D-E).

**Figure 3.**
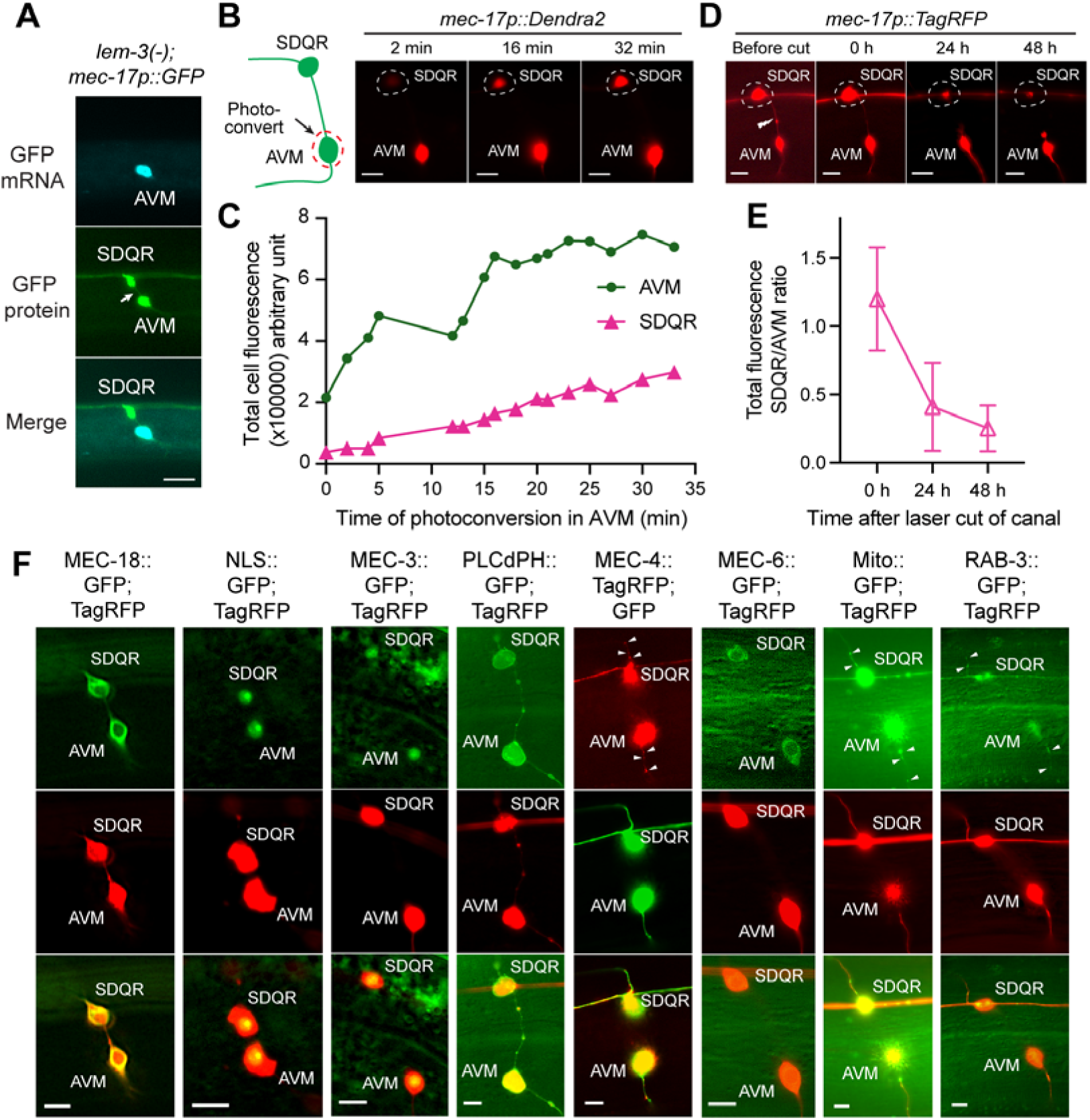
Cytoplasmic contents are mixed between the two cell bodies of the syncytium. (A) smFISH signals for the GFP mRNAs were only observed in the AVM cell body of the AVM-SDQR syncytium in the *lem-3(unk84)* animals carrying the *uIs31[mec-17p::GFP]* transgene. (B) A schematic cartoon and fluorescent images of the photoconversion experiments, in which dendra2 in the AVM was photoactivated by 488-nm lasers and the red fluorescence in SDQR was monitored over time. Scale bars = 5 μm. (C) Quantification of the converted dendra2 fluorescence in AVM and SDQR in the time course of photoconversion; a representative case is shown. (D) The change of RFP fluorescence in the SDQR cell body of the AVM-SDQR syncytium after the intercellular canal was severed. (E) The ratio of RFP fluorescent intensity of SDQR to that of AVM before and after the severing of the canal. (F) The distribution of various proteins fused with GFP or TagRFP (expressed from the *mec-17* promoter) in the AVM-SDQR syncytium. A diffusive TagRFP or GFP was co-expressed to allow the labelling of the syncytium. Arrows point to the signal in the neurites.

To assess what types of proteins can pass through the intercellular canal, we expressed fluorescently tagged cytoplasmic proteins (e.g., MEC-18::GFP), nuclear proteins (e.g., GFP::NLS, MEC-3::EGFP), membrane proteins (e.g., PLCdPH::GFP, MEC-4::TagRFP), ER-associated proteins (e.g., MEC-6::GFP), synaptic vesicle proteins (e.g., GFP::RAB-3), and mitochondrial proteins (e.g., mitochondrial matrix tagged GFP) from TRN promoters, which were activated in the AVM nucleus. The presence of these proteins in the SDQR cell body would indicate that they can pass through the canal. In all cases, we observed the fluorescent signals at comparable intensity in both cell bodies of the syncytium, indicating that diffusion (and/or transport) of the proteins through the canal led to even distribution in the two connected cells (Fig. 3F). Organelle-associated proteins maintained their normal subcellular localization in both cell bodies, suggesting that they could be potentially functional in both cells. Overall, the mixing of diffusible cytoplasmic content in the binucleate syncytium appeared to be thorough despite the largely distinct gene transcription profiles of the two nuclei.

### Unresolved chromatin bridges and connected nuclear envelopes lead to the formation of intercellular canal

Next, we investigated the cause of the intercellular canal connecting TRNs and their sister cells in *lem-3*(-) mutants. Since LEM-3 and its human ortholog ANKLE1 are required to enzymatically process chromatin bridges that arise from mitotic nondisjunction (*2, 5, 11*), we hypothesized that the intercellular canals connecting TRNs and their sister cells were induced by unresolved chromatin bridges in the *lem-3*(-) mutants. Since the penetrance of the TRN-sister syncytia was low (< 3% at 20°C or lower) in *lem-3*(-) mutants, we reasoned that the frequency of spontaneous chromosome segregation errors that need to be processed by LEM-3 is low. Interestingly, elevating the *C. elegans* culturing temperature to 25°C increased the frequency of AVM-SDQR and PLM-ALN syncytia (Fig. 4A), potentially due to the more frequent DNA replication and chromosomal segregation errors at higher temperature (*17–19*). In fact, disrupting DNA synthesis by knocking down the DNA replication licensing factor *mcm-7* (*2*) or treating the animals with the DNA topoisomerase II inhibitor ICRF-193 (*20*) significantly increased the penetrance of forming the syncytia in *lem-3*(-) mutants (Fig. 4A). Since ICRF-193 is known to induce chromatin bridges and sister chromatid nondisjunction in anaphase (*20, 21*), the result supports that the intercellular canal was formed due to unresolved chromosomal linkages. Nevertheless, elevated temperature and the ICRF-193 treatment had only marginal effects on increasing the penetrance of ALM-BDU and PVM-SDQL syncytia (Fig. 4B).

**Figure 4.**
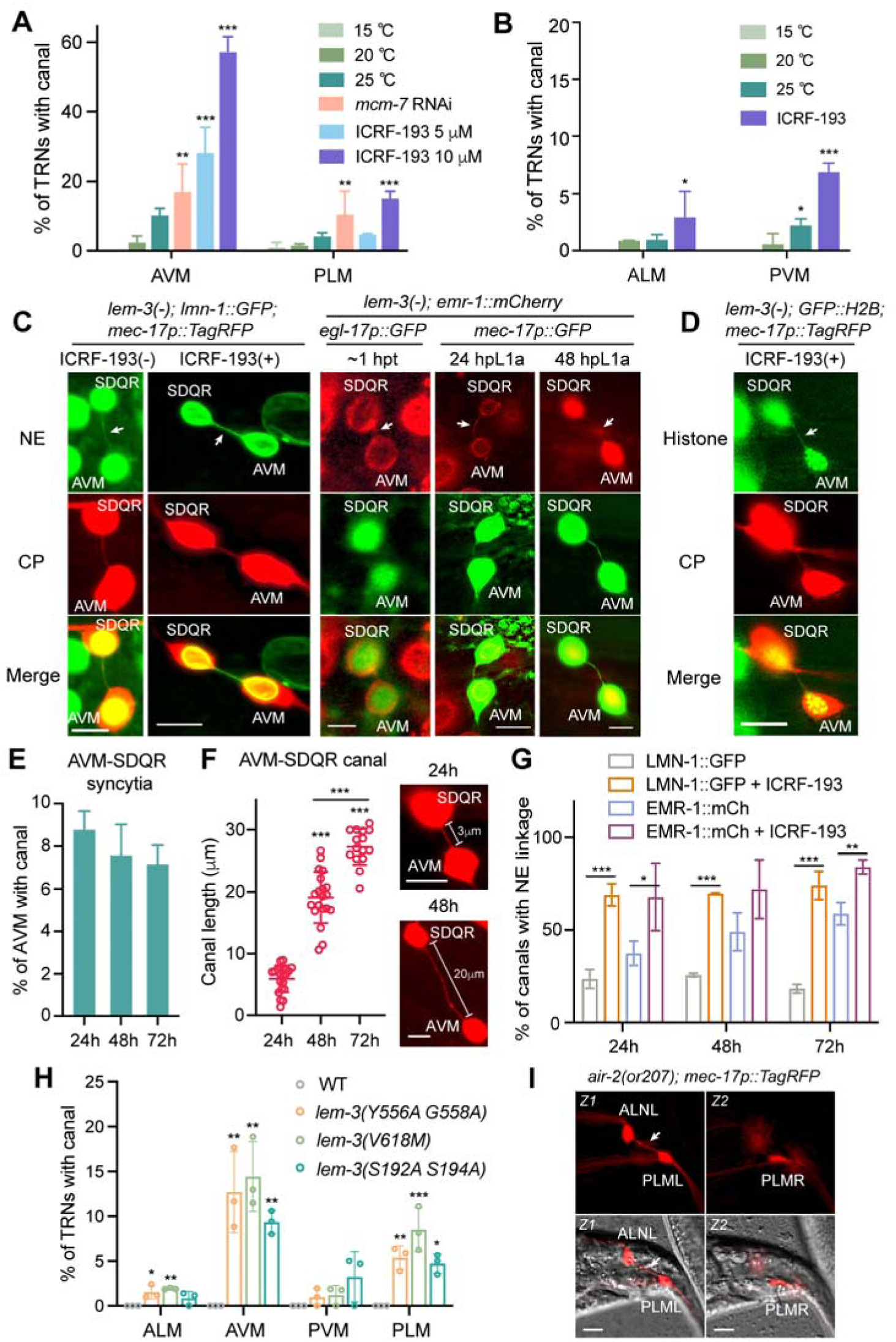
Unresolved chromatin bridges in *lem-3*(-) mutants led to the formation of the binucleate syncytium. (A-B) The penetrance of the TRN syncytia in *lem-3(unk84)* mutants grown under various conditions. Mean ± SD is shown. Single, double, and triple asterisks indicate *p* < 0.05, 0.01, and 0.001, respectively, in comparison with the 15°C condition in a Dunnet’s test. (C) Nuclear envelope (NE) is marked by the *ccIs4810[lmn-1p::lmn-1::GFP]* transgene in adult *lem-3(unk84)* mutants treated with or without 10 μM ICRF-193 in the left panel and by the *bqSi226[emr-1p::emr-1::mCherry]* transgene during development at 1 hour post telophase of the AVM-SDQR cell division (hpt) and 24 and 48 hours post L1 arrest recovery (hpL1a). Another transgene of different color was used to label the TRN syncytium. Arrows point to the NE linkage. Scale bars = 5 μm. (D) Histone is labelled using the transgene *oxTi75[eft-3p::GFP::H2B]* and is found in the intercellular canal (arrow). (E) Percentages of animals showing AVM-SDQR syncytia with an intercellular canal at different time post L1 arrest recovery (left panel); percentages of syncytia showing NE linkages with two different markers in the presence and absence of 10 μM ICRF-193 (right panel). Asterisks indicate significance in a *t*-test. (F) The length of the intercellular canal of the AVM-SDQR syncytia at different time post L1 arrest recovery. (G) Penetrance for the TRN syncytia in *GFP::lem-3* missense *gt3311(Y556A G558A)* and *gt3310(S192A S194A)*, and *lem-3(unk14; V618M)*. Significance in a Dunnett’s test comparing the mutant with the wild type is shown. (H) PLML-ALNL syncytium in *air-2(or207ts)* mutants grown at 25°C from early embryonic stages. “Z1” and “Z2” mean two focal planes of the same animal.

To confirm the presence of chromatin bridges, we found histones (H2B) and nuclear envelope markers (e.g., LMN-1/Lamin (*22*) and EMR-1/Emerin (*23*)) in the canal (Fig. 4C-D), although attempts to directly visualize DNA using various dyes were not successful. Importantly, these bridges (e.g., between AVM and SDQR nuclei) were formed immediately after the telophase of the scheduled cell division and can last for several days throughout development and into the adulthood of the animal. When treating the animals with ICRF-193, we observed much stronger signal for nuclear envelope markers in the canal (Fig. 4C). Nevertheless, only ∼20% of AVM-SDQR syncytia showed persistent nuclear envelope connection 16 hours after the cell division (i.e., 24 hours after recovery from L1 arrest), suggesting that the DNA bridge could break as the two nuclei migrated away from each other during development (Fig. 4E).

In fact, we observed the elongation of the AVM-SDQR canal from ∼6 μm in average length in L1/L2 animals to ∼27 μm in adults, and the ALM-BDU canal can extend to >100 μm in length (Fig. 4F). The stretching of the chromatin bridge may have led to the mechanical breakage and the loss of nuclear linkage, but the cytoplasmic linkage through the canal (a membrane structure) was maintained likely due to the inactivation of cytokinesis machineries (Fig. 4E). Compared to the spontaneously formed chromatin bridges, ICRF-193-induced chromatin bridges appeared to be more resistant to mechanical breakage likely due to the accumulated chromosomal aberration, since significantly more syncytia maintained the nuclear membrane connection in adults under ICRF-193 treatment compared to untreated *lem-3*(-) mutants (Fig. 4G). When quantifying the distance between the AVM and SDQR nuclei, we found that some syncytia with long nuclear distance could still maintain the nuclear envelope connection into adulthood in the absence of ICRF-193 treatment, suggesting that some naturally occurring DNA linkages might be strong enough to survive the stretching (Fig. S8A-B). Interestingly, we also found some rare cases where the two nuclei stayed in the same cell body (thus, there is not canal); we reason that either furrow ingression never occurred, or regression happened due to the presence of chromatin bridges in *lem-3*(-) mutants (Fig. S5D).

The endonuclease activity of LEM-3 is required to cleave the unresolved chromatin bridges during early embryogenesis (*2*). As expected, the Y556A and G558A double mutations in the conserved GIY-YIG nuclease motif caused the formation of TRN-sister cell syncytia at similar penetrance as the *lem-3*(-) deletion allele (Fig. 4G). Interestingly, the V618M mutation found in the *unk14* allele we isolated from the genetic screen was also mapped to the nuclease domain. Moreover, Aurora B kinase-mediated phosphorylation of LEM-3 is required for its activity in resolving the bridge and preventing the syncytium, because the S192A and S194A double mutation of the putative Aurora B phosphorylation sites (*2*) caused phenotypes similar to *lem-3*(-) mutants. Inactivation of *air-2*, which codes for Aurora B in *C. elegans*, also led to the formation of the syncytia connecting TRNs and their sister cells, but we only observed the PLM-ALN syncytia using a temperature-sensitive (and likely partial loss-of-function) *air-2* allele (Fig. 4H).

In addition to LEM-3, we also examined the effects of other endonucleases, including MUS-81/MUS81, SLX-1/SLX1, and XPF-1/ERCC4, which are involved in resolving Holiday junctions, DNA replication errors, and interstrand crosslink, as well as DNA repair (*24–26*). Nevertheless, we did not observe any TRN-sister syncytium in either *xpf-1*(-) single mutants or *xpf-1*(-)*; mus-81*(-) double mutants or *mus-81*(-) *slx-1*(-) mutants, suggesting that LEM-3 may have specific functions in cutting chromatin bridges connecting sister cells.

### The intercellular canal contains cytoskeleton structures

When chromatin bridge forms, LEM-3 is recruited to the midbody, which is a transient microtubule structure formed within the intercellular connection between sister cells prior to abscission (*2*). Using the centralspindlin protein ZEN-4/MKLP1 as a midbody marker (*27*), we found that midbody formed normally in *lem-3*(-) mutants and persisted in the intercellular canal for hours likely because the unresolved chromatin bridges aborted cytokinesis (Fig. 5A-B). However, the ZEN-4 signal eventually disappeared 6 hours after the telophase, suggesting that midbody structure cannot last for a long time if abscission does not occur (Fig. 5B).

**Figure 5.**
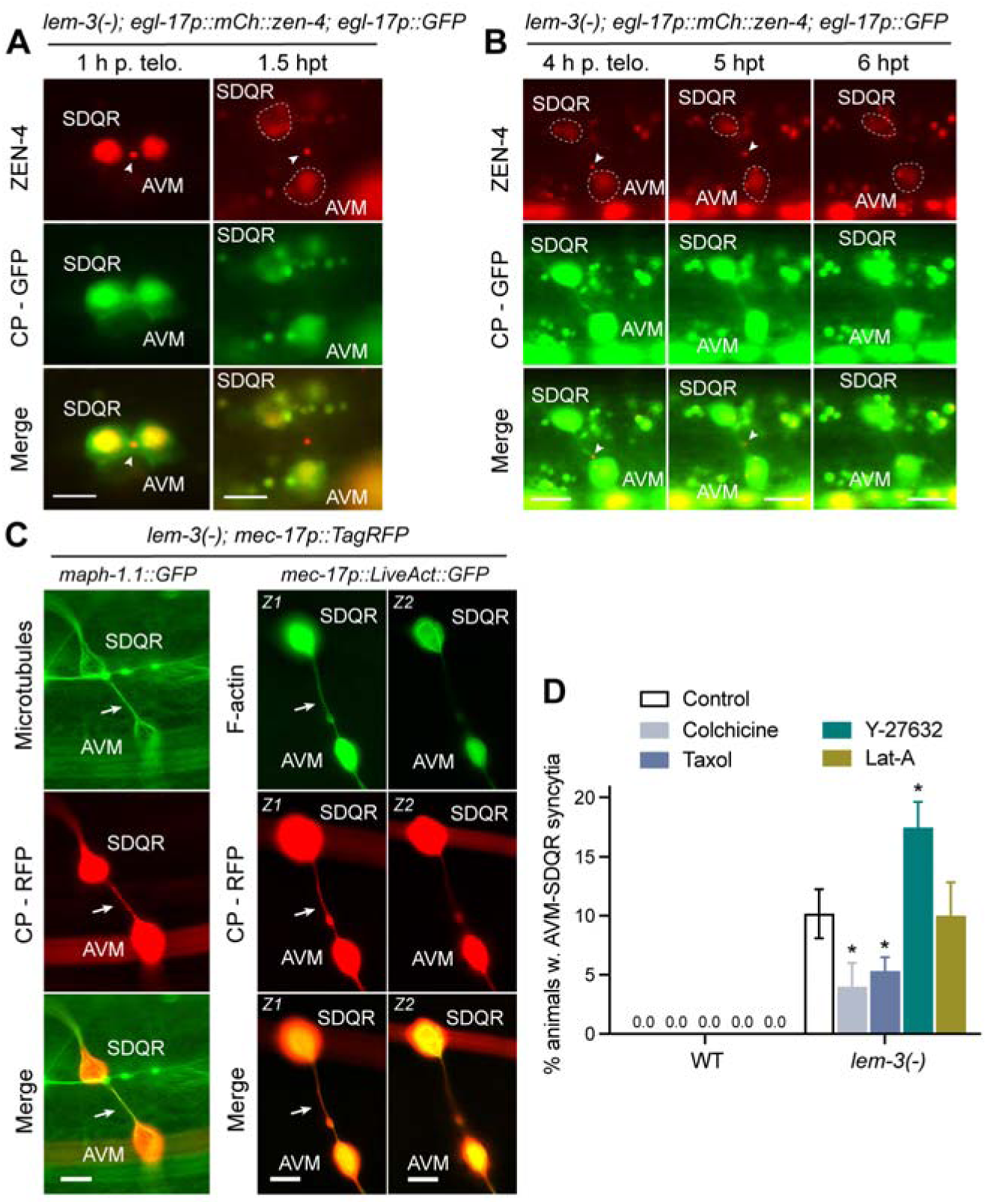
Failed cytokinesis leads to the formation of intercellular canal which is stabilized by microtubules. (A) The midbody complex labelled by mCherry::ZEN-4 expressed from the *casIs131* transgene in the AVM-SDQR syncytium at 1- and 1.5-hours post telophase (hpt) of the cell division. Dashed lines indicate the cell bodies, as cytoplasm (CP) is labelled by GFP expressed from *ayIs9[egl-17p::GFP]*. Arrow heads point to the midbody. Scale bars = 5 μm. (B) Tracking of the mCherry::ZEN-4 signal in the same syncytium shows the disappearance of the midbody complex at 6 hpt. (C) Microtubules and F-actin in the intercellular canal of the AVM-SDQR syncytium is labelled by MAPH-1.1::GFP (MAPH1.1 is a MAP1 family protein) expressed from *unkEx170[maph-1.1p::maph-1.1::GFP]* and LiveAct::GFP expressed from *unkEx572[mec-17p::LiveAct::GFP]*, respectively. Arrows indicate the canal. “Z1” and “Z2” mean two focal planes of the same animal. (D) The percentage (mean ± SD) of wild-type (WT) and *lem-3(unk84)* animals showing the AVM-SDQR syncytia when treated with 1 mM colchicine, 100 nM taxol, 100 nM Y-27632 (ROCK inhibitor), or 1 μM Latrunculin-A (Lat-A). Single asterisk indicates *p* < 0.05 in comparison with the untreated control in a Dunnett’s test.

On the other hand, by visualizing the microtubules using a GFP-fused microtubule-associated protein MAPH-1.1 (a homolog of human MAP1 proteins) (*28*), we found that microtubule structure persisted in the intercellular canal and likely provided structural support for the formation and maintenance of the canal (Fig. 5C). Interestingly, treatment with both microtubule-destabilizing and -stabilizing drugs (colchicine and paclitaxel, respectively) reduced the penetrance of AVM-SDQR syncytia, indicating that optimal microtubule stability and dynamics are crucial for forming or maintaining the intercellular canal. With the structural support provided by microtubules, the canal could persist into even aged animals (e.g., day 5 adults; Fig. S8C).

Using *LifeAct::GFP* that binds to actin filaments (*29*), we also found actin filaments in the intercellular canal (Fig. 5C). The presence of both microtubule and actin cytoskeletons suggested that the canal may be maintained as a regular neuronal process by the syncytium. However, treating the animals with Latrunculin A that blocks actin polymerization did not affect the penetrance of AVM-SDQR syncytia, suggesting that the formation and maintenance of the canal may not require actin filaments (Fig. 5D). On the contrary, inhibiting ROCK-induced myosin phosphorylation using the ROCK inhibitor Y-27632, which is known to delay furrow ingression (*30*), significantly increased the penetrance of forming the AVM-SDQR canal in *lem-3*(-) mutants (Fig. 5D). This finding supports that unresolved chromatin bridges and weaker actomyosin contractility would delay cytokinesis and eventually abort abscission, resulting in permanent intercellular canals between postmitotic sister cells.

### Connection between apoptotic and non-apoptotic cells in *lem-3*(-) mutants

When examining the AVM-SDQR syncytia, we noticed that ∼25% of them had a third cell body, which was connected with the other two cell bodies through intercellular canals (Fig. 6B). We referred to these structures as “three-body syncytia”. Since the sister cell (QR.pp) of the precursor (QR.pa) that generates AVM and SDQR undergoes apoptosis (Fig. 6A) (*31*), we reasoned that this additional cell body may be the remains of the apoptotic cell. This third cell body contained a nucleus labelled by the nuclear envelope marker, was generated through cell division (indicated by the presence of the midbody), but did not transcribe TRN genes (Fig. 6B-D). Interestingly, as the animal develops, the frequency of the three-body syncytia decreased over time (Fig. 6E), implying that the third cell body could be removed (likely through engulfment by other cells) without inducing programmed cell death of the entire syncytium. This finding also suggests that the apoptotic fate of the precursor’s sister (QR.pp) may only be delayed but not reversed through the connection with the surviving AVM and SDQR neurons.

**Figure 6.**
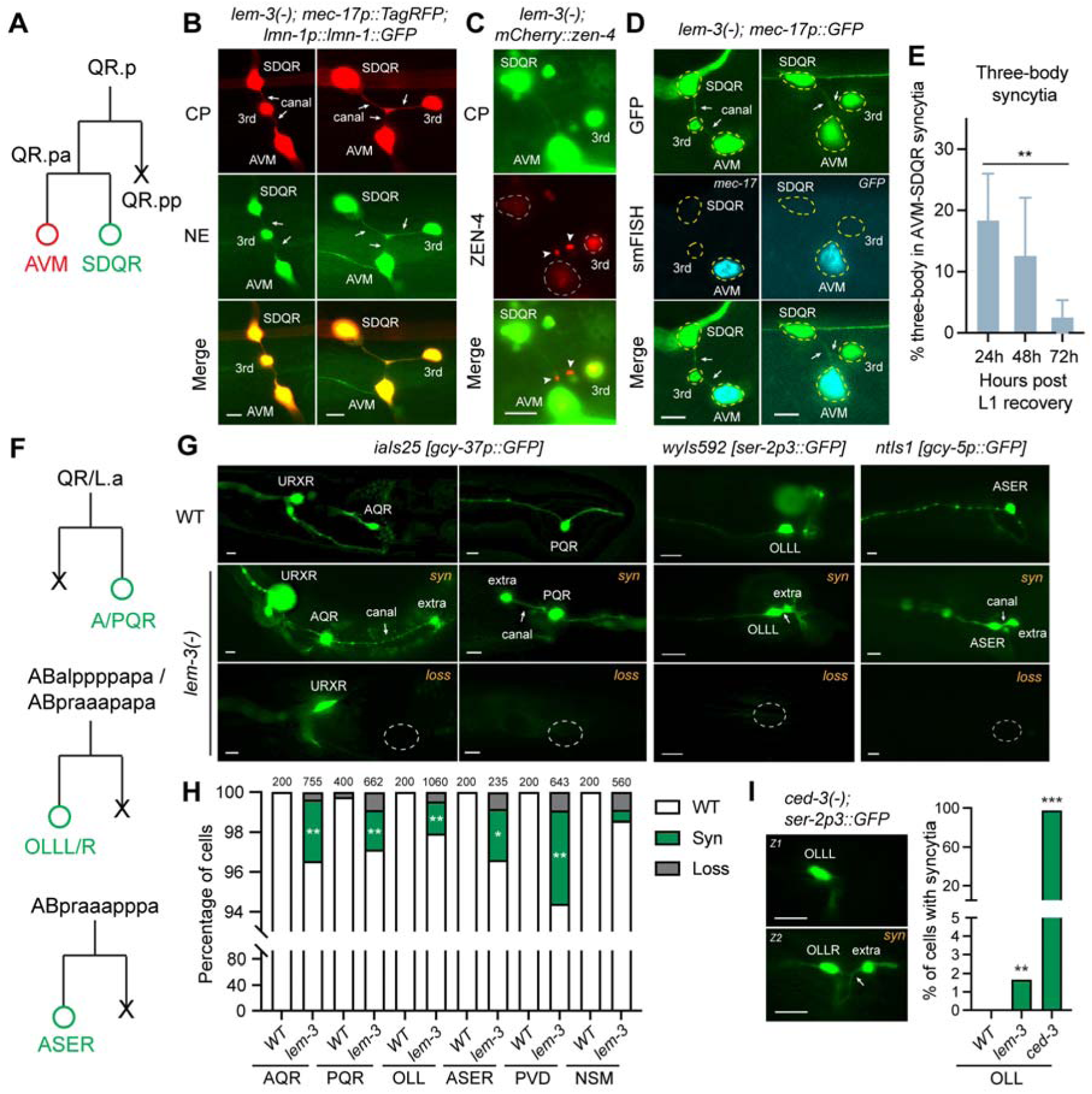
Binucleate syncytium affects the apoptotic fate of the sister cells. (A) The cell lineage that generates the precursor of AVM and SDQR and an apoptotic sister (“X” means apoptosis). (B) Three-body syncytia connecting SDQR, AVM, and a third cell body (“3rd”), connected by intercellular canals (arrows), which contain nuclear envelope (NE) links labelled by LMN-1::GFP. Scale bars = 5 μm. (C) The canal contains two midbody complexes labelled by mCherry:ZEN-4 (arrow heads) from two rounds of cell divisions that result in the three-body syncytium. (D) smFISH signal for endogenous *mec-17* and exogenous GFP in the three-body syncytium of the *lem-3(unk84); uIs31[mec-17p::GFP]* animals. (E) The percentage of AVM-SDQR syncytia that possess a third body throughout development. (F) The cell lineages that give rise to A/PQR, OLLL/R, and ASER neurons and their apoptotic sisters. (G) Representative images of the two phenotypes were observed for the three types of neurons in *lem-3(unk84)* mutants: the syncytia formed between them and their apoptotic sisters (“*syn*” and extra cell body) and the loss of the fate marker labelling (“*loss*” and dashed circle). (H) Penetrance of the “*syn*” and “*loss*” phenotypes for the indicated neuron. Single and double asterisks indicate *p* < 0.05 and 0.01, respectively, in a Chi-square test comparing the frequencies of the “*syn*” phenotype between WT and *lem-3*(-) animals. The number of cells examined is shown above the stacked bar. (I) Representative images of *ced-3(n717); wyIs592[ser-2p3::GFP]* animals showing the syncytium phenotype; quantification of the penetrance is shown on the right (significance from a Dunnett’s test is shown).

The above observation prompted us to examine cell divisions that generate one terminally differentiated neurons and one apoptotic cell. Among the four cell types we examined, AQR and PQR neurons form syncytia with their apoptotic sisters at detectable penetrance (2-3%) through oftentimes a long intercellular canal (especially for AQR) (Fig. 6F-G), which may be due to the migratory nature of these neurons (*31*). We also observed some rare cases where the AQR and PQR neurons were missing (they likely underwent apoptosis or failed to differentiate), suggesting that neuronal survival and differentiation could also be affected by cytoplasmic connection with cells that were scheduled to die. This may be similar to how cytokinetic bridges in mouse embryos allow the sharing of apoptotic factors to coordinate cell elimination (*32*). We did not observe such loss of marker expression in the AVM-SDQR syncytia, likely because only their precursor is connected to an apoptotic sister and thus the effect might be weaker.

Similar results were obtained for OLL, ASER, and PVD neurons with the penetrance of syncytia ranging from 1.6% to 4.7% and the penetrance of cell fate marker loss below 1% (Fig. 6F-H and S9). It appears that connecting a postmitotic neuron with its apoptotic sister is more likely to save the sister cell from dying than to induce the apoptosis (or failure in differentiation) of the entire syncytium. No statistically significant phenotype was observed for NSM neurons. To further confirm that the extra cell is caused by blocked apoptosis, we crossed the OLL fate marker with *ced-3*(-) mutants (*ced-3* codes for caspase 3) and found that >95% of the OLLs were connected with an extra cell in the mutants (Fig. 6I). The low penetrance of the phenotype in *lem-3*(-) mutants supports that chromatin bridges between two sister cells are generally rare.

### LEM-3 safeguards the specification of many neuron types

To understand whether LEM-3 has a general function in preventing the formation of syncytia and safeguarding cell fate specification in the nervous system, we next extended our analysis to eleven other cell divisions that generate two daughter cells with distinct neuronal fates (including the CEPD-URX, SMDD-AIY, SMDV-BAG, ASJ-AUA, ADF-AWB, HSN-PHB, AFD-RMD, ADE-ADA, PDE-PVD precursor, FLP-AIZ, and RIS-DB4 sister pairs). We used fluorescent reporters to mark one of the two cell fates in a sister pair in the *lem-3*(-) mutants. Interestingly, in almost all cases, we observed both the formation of syncytium with an additional cell body (presumably the sister cell) and the loss of the marker expression (indicating fate suppression likely due to the connection with the sister cell) with one being the dominant phenotype despite the low penetrance (Figure 7A-F and S10). One exception is the RIS neuron, where we did not observe any significant phenotype in the *lem-3*(-) mutants, but ICRF-193 treatment led to ∼20% of syncytium formation (Figure 7F and S10F). We also used the CEPD-URX sister pair as an example to confirm that the appearance of syncytium labelled by the URX marker (*gcy-37p::GFP*) coincided with the loss of the CEPD marker (*dat-1p::mCherry*) in the same animal (Figure 7C). In fact, the penetrance of syncytium for the URX marker was similar to the penetrance of fate loss with the CEPD marker.

**Figure 7.**
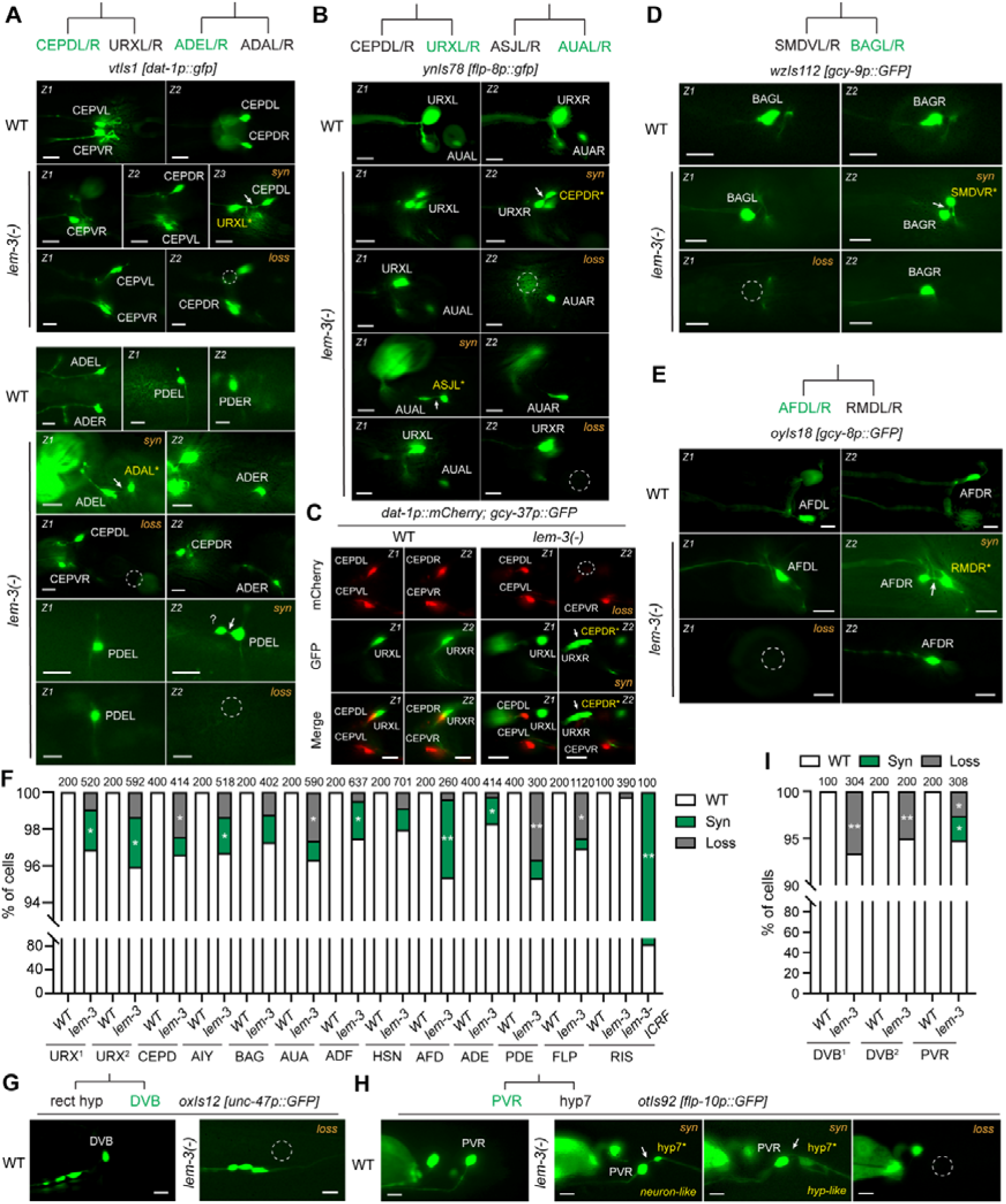
Mutations in *lem-3* lead to the formation of syncytia for many sister cell pairs. (A) Dopaminergic neurons CEPDL/R, CEPVL/R, ADEL/R, and PDEL/R are labelled by *dat-1p::GFP* and showed both syncytium (“*syn*”) and fate loss (“*loss*” and dashed circle) phenotype in the *lem-3(unk84)* mutants. For the syncytia, the abnormally labelled sister cells have their names in yellow; arrows indicate the intercellular canal. It should be noted that PDE’s sister is the precursor of PVD and its apoptotic sister (see Figure S9A). Scale bars = 5 μm. “Z1” and “Z2” mean two focal planes of the same animal. (B) URXL/R and AUAL/R are labelled by *ynIs78[flp-8p::GFP]* and showed the syncytium and fate loss phenotypes. (C) The CEPD and URX fates are labelled by *dat-1p::mCherry* and *flp-8p::GFP*, respectively, in the same *lem-3*(-) animal. The fate loss of CEPDR coincide with the URX syncytium. (D-E) Representative images for the syncytium and fate loss phenotypes for BAGL/R and AFDL/R neurons in *lem-3*(-) mutants. (F) Penetrance of the “*syn*” and “*loss*” phenotypes for the indicated neuron. Single and double asterisks indicate *p* < 0.05 and 0.01, respectively, in a Chi-square test comparing the frequencies of the indicated phenotype between WT and *lem-3*(-) animals. The number of cells examined is shown above the stacked bar. URX^1^ indicates the results of using *iaIs25[gcy-37p::GFP]*, and URX^2^, the results of *ynIs78[flp-8p::GFP]*. (G) The loss (dashed circle) of DVB fate marker in *lem-3*(-) mutants. (H) The gain of extra (neuron-like and hypodermis-like) cells labelled by PVR marker and the loss of PVR fate marker in *lem-3*(-) mutants. (I) The penetrance of the “*syn*” and “*loss*” phenotypes for DVB and PVR neurons. DVB^1^ indicates the results of using *oxIs12[unc-48p::GFP]*, and DVB^2^, the results of *otIs92[flp-10p::GFP]*.

Lastly, since neurogenesis in *C. elegans* is largely non-clonal, we also analysed terminal cell divisions that happen to generate one neuron and one non-neuronal sister. For example, the sister cell of the DVB neuron is the rectal cell, and we found that the expression of two DVB markers (*unc-47p::GFP* and *flp-10p::GFP*) was both lost in ∼5% of the *lem-3*(-) animals, suggesting that the DVB fate may potentially be suppressed in the syncytium (Figure 7G and 7I). In the case of PVR neuron, whose sister is the hypodermal cell hyp 7, we found both syncytium and fate loss phenotypes with equal penetrance in *lem-3*(-) mutants. Interestingly, the extra cell connected to PVR in the syncytium appeared to be neuron-like in some animals and hypodermis-like in others based on the morphology (Figure 7H-I). This finding hinted that it may be possible to convert skin cells into neurons through direct cytoplasmic coupling, although this conversion was previously demonstrated by overexpressing certain transcription factors (*33*) or microRNAs (*34*). On the other hand, just like the AVM-SDQR syncytium, two cell fates that are as distinct as neuron and skin might co-exist in the same syncytium that links the two cell bodies through an intercellular canal.

## Discussion

In this study we uncovered a role of the endonuclease LEM-3 in the separation of sister cells during final round of cell division prior to terminal differentiation and fate specification. The loss of LEM-3 led to unresolved chromatin bridges which resulted in cytokinesis failure and formation of binucleate syncytia. Based on the separation of the two nuclei, these syncytia exhibit cellular ploidy rather than nuclear ploidy, suggesting successful karyokinesis but failed cytokinesis. This phenotype is consistent with the established function of LEM-3 as a midbody-tethered nuclease that resolves persistent DNA bridges, acting as a “last-chance” mechanism to ensure proper cell abscission (*12, 35*). However, syncytia found in *lem-3*(-) mutants are mostly “two-body” syncytia, which have two distinct cell bodies that are connected by an intercellular canal, instead of the commonly observed “one-body” syncytia, where furrow regression leads to a big cell body that contains two nuclei. Although many factors (e.g., mutations in cytoskeleton regulators, loss of DNA repair factors, kinetochore defects, chromatin linkages) can cause cytokinesis failure and polyploidy (*36, 37*), such “two-body” syncytia were rarely observed. This is likely because most of the previous reports studied cell division using continuously dividing cells (either cells in culture or embryonic cells), whereas we studied the final round of cell division in development. The persistent chromatin bridges in these postmitotic cells appear to neither evoke mechanical resolution nor cause furrow regression. The pause of abscission triggered by the Aurora B-dependent NoCut checkpoint became a permanent state for the two sister cells; then the stretch of the chromatin bridge as the two nuclei move away from each other leads to the formation of the intercellular canal and the “two-body” syncytia. Although the underlying mechanisms for such phenomenon is not entirely clear, we reason that it may involve the disassembly of the midbody structure on the canal and the exit of cell cycle.

The two nuclei in the *lem-3*(-) syncytia (e.g., AVM-SDQR syncytium) could maintain distinct transcriptional programs despite the nuclear and cytoplasmic connection. This finding is consistent with the results of single-nucleus RNA sequencing and spatial transcriptomic studies on various types of syncytia (including skeletal muscle fibers, placental trophoblasts, and slime mold plasmodia), which also revealed transcriptional heterogeneity driven by positional cues, developmental stages, or environmental stimuli (*38–41*). However, the functional consequences at the protein level of this transcriptional heterogeneity is not entirely clear. Our work offers an example to show that proteins translated from the transcripts synthesized in one nuclei can easily diffuse into the entire syncytium with a size of a few hundred micrometers in length. At this scale, the transcriptional diversity of the two nuclei does not lead to protein diversity between the two connected cell bodies. At a larger scale (e.g., muscle fibers can reach tens of centimeters in length), how much protein heterogeneity exists in a single syncytium is unclear but can be understood by detailed spatial proteomic studies with subcellular resolution.

Multinucleated syncytia, in physiological context, can promote the functionality of tissues, such as the heart muscles (cardiomyocyte syncytia), bone marrow (megakaryocytes), and liver (hepatocyte syncytia), by enhancing metabolic capacity and stress resistance (*42*). Under pathological conditions, however, polyploidy is linked to blood disorders, female infertility, age-related macular degeneration, and cancers (*36, 37*). The loss of human ANKLE1 impaired erythrocyte differentiation by the failure in cleaving and degrading mitochondrial DNA (*43*) and led to polyploidy that renders breast cancer cells in a stem cell-like state (*44*). Interestingly, the binucleate syncytia in the *lem-3*(-) mutants in our study did not revert to a precursor state despite the mixing of the cellular contents of the two sister cells. Instead, in some case (e.g., the AVM-SDQR syncytium), the two connected cells were able to undergo morphological and molecular differentiation to adopt their respective cell shapes and activate a battery of genes associated with their respective fates. This developmental robustness against polyploidy is surprising and suggests that two independent modules control the differentiation of two neuron types (e.g., AVM and SDQR). They do not interfere with each other and can essentially operate in parallel in the syncytium to allow it to develop into a mixed fate.

On the other hand, the ALM-BDU and PLM-ALN syncytia represent a different type of examples, where the fate of one of the sister cells is suppressed and the two nuclei in the syncytium adopt similar transcriptional programs (i.e., the TRN genetic program). A potential reason for the dominance of the TRN fate in the syncytia is that the TRN terminal selector MEC-3 can diffuse into the sister cells through the intercellular canal and exert an dominant effect to activate the TRN fate and repress the sister cell fate. A previous study showed that MEC-3 has higher affinity to the transcription factor UNC-86 than the BDU fate selector PAG-3 (*14*). Thus, the presence of MEC-3 in the BDU not only causes MEC-3/UNC-86 heterodimer activating TRN genes but also prevents the formation of PAG-3/UNC-86 complex required for the activation of BDU genes. Interestingly, the BDU cell in the ALM-BDU syncytia is only partially converted to the ALM fate (e.g., the expression of BDU marker *ceh-14* is maintained) likely due to the moderate level of MEC-3 in BDU. Ectopic overexpression of MEC-3 in BDU can fully transform the morphology and gene expression of BDU into a ALM-like state.

The loss of BDU and ALN fates in *lem-3*(-) mutants suggests a detrimental effect of cytoplasmic linkage between two sister cells. In fact, many fate specification processes we examined are affected (i.e., loss of marker expression) by mutations in *lem-3*, although at a low penetrance due to the low frequency of naturally occurring chromatin bridges. Thus, we suspect that many fates are likely vulnerable to a competing fate selector from the sister cell. In this sense, by resolving the chromatin bridges, LEM-3 prevents the formation of binucleate syncytia and safeguard the fate specification process. In addition, *lem-3*(-) mutants can serve as a tool to study the interaction between the fate-determining programs of two sister cells (e.g., coexistence or dominance of one against another). Given that LEM-3 is an evolutionarily conserved endonuclease and has conserved functions in processing DNA bridges, future work can explore whether its orthologs also play important roles in terminal differentiation in other organisms.

## Materials and Methods

### Strains and constructs

*C. elega*ns strains were maintained at 20°C on nematode growth medium (NGM) plates seeded with OP50 bacteria according to previous methods (*45*). Unless otherwise indicated, the phenotypes of *lem-3* mutants were scored at 25°C, which produced the highest penetrance among the conditions we tested. To generate the *lem-3* knockout, we used CRISPR/Cas9-mediated gene editing to make cuts at two sites of the endogenous locus. The target sites 5’-GTGGTGATTGCTCCGTTTGG-3’ in exon 1 and 5’-AACAATCGACGTGGAGCAGG-3’ in exon 7. We synthesized the single guide RNAs using the EnGen sgRNA Synthesis Kit (E3322V) from NEB and injected them together with 20 pmol recombinant Cas9 (EnGen S. pyogenes Cas9 NLS from NEB, M0646T) into *C. elegans* using pCFJ104 (*myo-3p::mCherry*) as a co-injection marker. We genotyped the transformants with red muscles to screen for deletions in *lem-3* and obtained *unk84* and *unk85* alleles. Both alleles deleted the sequence between the two target sites, while *unk84* had a short insertion of 24-bp random sequences (Fig. 1A). We used *unk84* as the *lem-3*(-) mutant in most experiments. The *gt3309(eGFP::Stag::lem-3)*, *gt3310(eGFP::STag::lem-3[S192A S194A])*, and *gt3311(eGFP::Stag::lem-3[Y556A G558A])* alleles (*2*) were generously provided by Dr. Anton Gartner and Dr. Ye Hong. To rescue *lem-3*(-) mutant phenotype, we cloned a 9-kb *lem-3* genomic fragment, including 6 kb sequences upstream of the start codon, the coding region, and the 3’UTR, into pUC57 and injected it into the *lem-3(unk84)* mutants. To label various neuronal fates, we used an array of transgenic fluorescent reporters generated by previous studies and crossed them with the *lem-3* mutants. All strains used in this study is listed in Table S1.

To misexpress *mec-3*, we cloned a 2,188-bp *unc-119* promoter between HindIII and BamHI sites in pPD95.75 and then inserted the *mec-3* coding and 3’UTR sequence at the NotI site. The resulted construct was then injected into *C. elegans* gonad and integrated into the genome to form *uIs158[unc-119p::mec-3]*. To create transgenes that express proteins with various subcellular localizations under the TRN-specific promoters (e.g., a 1.9-kb *mec-17* and a 400-bp *mec-18* promoters), we cloned the respective proteins (e.g., dendra2 for photoconvertible fluorescent protein, the PH domain of PLCdelta1 fused with GFP for membrane protein (*46*), GFP fusion protein with a nuclear localization signal, the ER chaperone MEC-6 for ER (*47*), etc.; see Table S1 for details). To label microtubules, we cloned a genomic fragment of *maph-1.1* (a homolog for MAP1), including a 3.7-kb promoter, coding region and 1-kb 3’UTR and inserted GFP before the stop codon. To label actin filaments, we fused the LiveAct sequence (*29*) at the N-terminus of GFP and inserted it downstream of the *mec-17* promoter. These constructs were injected into *C. elegans* gonad to form extrachromosomal arrays, some of which were integrated using gamma irradiation.

### smFISH and smiFISH

To monitor transcription of endogenous genes from the two nuclei of the syncytia, we perform single-molecule RNA fluorescent in situ hybridization (smFISH) against TRN-specific genes *mec-17*, *mec-18*, and *mec-3*, and the sister fate-specific genes *lad-2*, *zig-3*, *ast-1*, and *D2089.3*. To detect transcription of GFP in the syncytia that express the integrated fluorescent fate reporters, we conducted smFISH against GFP. In general, we designed the FISH probes using the Stellaris RNA FISH Probe Designer tool from Biosearch Technologies (Petaluma, CA) and synthesized them with Quasar 670 dye. L1 or L2 stage animals from various strains were fixed, stained with the FISH Probe sets in the hybridization buffer (purchased from Biosearch Technologies), and then washed using the wash buffer based on a previous protocol (*48*). The hybridized animals were then mounted using VECTASHIELD Antifade Mounting Medium (Vector Laboratories #H-1000) and imaged using a Leica DM8 microscope with Cy5 filter sets.

We also performed smiFISH (single-molecule inexpensive FISH) (*49*) against additional TRN sister fate-specific genes, including *nlp-20*, *nlp-80*, *flp-12*, and *C15C7.4*. smiFISH is a modified version of smFISH in which the primary probe binds to the target mRNA and contains a flap sequence that is bound by a secondary probe which is fluorescently labelled. We designed the primary probes using the Stellaris RNA FISH Probe Designer tool and added the X Flap sequence to the 5’ end of each probe. The X Flap sequence was 5’-CCTCCTAAGTTTCGAGCTGGACTCAGTG-3’. An equimolar mixture of the primary probe set was prepared at 100 μM in IDTE buffer (10 mM Tris-HCl, 0.1 mM EDTA, pH 8.0). Secondary probes were the reverse complement of the X Flap sequence and were labelled with Quasar 670 dye at the 5’ end. Both primary and secondary probes were synthesized by Beijing Tsingke Biotech Company Limited. To hybridize the fluorophore-labelled secondary probes to the 5’-extended primary probe sets, an annealing reaction was performed at a final concentration of 20 _μ_M. The reaction mix consisted of 2 μl of 100 μM primary probe mix (200 pmol in total), 2.5 μl of 100 μM secondary probe (250 pmol in total), 1 μl of 10X T4 DNA ligase buffer (NEB), and 4.5 μl of water. The reaction mix was incubated in a thermocycler using the following programs: 95°C for 3 min, decrease 0.5°C per cycle and hold for 30 s (59 cycles), 65°C for 3 min, decrease 0.5°C per cycle and hold for 30 s (79 cycles), 95°C for 5 min. Steps for *C. elegans* sample preparation, hybridization, washing, and image acquisition were the same as those for smFISH. All FISH probes used in this study are listed in Table S2.

### Pharmacological treatments and *mcm-7* RNAi

To treat the animals with various drugs, we added ICRF-193 into the NGM plates at 10 μM, Colchicine at 1 mM, Paclitaxel (Taxol) at 100 nM, Y-27632 at 100 nM, and Latrunculin-A at 1 μM.

For *mcm-7* RNAi, we obtained the bacteria expressing dsRNA against *mcm-7* from the Ahringer *C. elegans* RNAi feeding library and grew the bacteria at 37°C until reaching an OD600 of 0.6 and then seeded the bacteria onto NGM plates supplemented with 1 mM IPTG for overnight induction. L4 hermaphrodites were transferred to the RNAi plates and fed at 25°C. Treating the animals with 100% *mcm-7* RNAi bacteria led to sterility, so we mixed *mcm-7* RNAi bacteria and OP50 of equal OD600 at 7:93 ratio to make a 7% *mcm-7* RNAi bacteria solution and seeded them on the RNAi plates.

### Photoactivation and photoablation

Day-one adult worms were immobilized on 10% agarose pads using polystyrene microspheres (Polysciences #00876–15; diluted 6-fold) and high-precision cover glasses (Paul Marienfeld GmbH & Co. KG #0107052). Worms were raised in the dark to avoid unnecessary photoconversion under natural lighting. As a small amount of Dendra2 is activated during worm handling, a Rhodamine fluorescence filter set was used to locate the TRN syncytium by detecting very low levels of red fluorescence. Using the green (546 nm) light to track the TRN syncytium served two purposes: 1) to avoid further depletion of the inactive Dendra2 pool, and 2) to photobleach activated Dendra2 as much as possible before photoconversion. A 405 nm laser was then used to deliver 2s of blue light at 5% intensity to the TRN cell bodies via a Leica Infinity Scanner powered by a Widefield Compact Supply Unit on a Leica DMi8 microscope equipped with 63x water objectives. The same 405 nm laser was applied to the TRN cell bodies each time when the red fluorescence in the connected sister cells was imaged using 10% intensity of 546 nm light at various intervals. The fluorescent intensity of activated dendra2 was quantified by using imageJ.

To cut the canal using laser, day-one adult animals were similarly immobilized, and 355-nm UV light was delivered to the middle of the canal through a using a Pulsed Laser Unit attached to the Infinity Scanner on the Leica DMi8 using 63x water lenses. To minimize injury, the lowest energy that enabled the severing of the canal was used. Animals were then rescued and placed on NGM plates for recovery and were imaged 24 and 48 hours after the axotomy. The fluorescence in both AVM and SDQR cell bodies were quantified.

### Image acquisition, phenotype scoring, and statistical analysis

Fluorescent imaging was conducted using a Leica DMi8 inverted microscope outfitted with a Leica K5 monochrome camera. The Leica THUNDER deconvolution system was applied to enhance image clarity and eliminate out-of-focus light in certain images. Image analysis and measurements were carried out using the Leica Application Suite X software. To score the penetrance of syncytium formation, animals were grown at 25°C for at least two generations and then were imaged at day-one adult stage. At least 200 animals from each strain were examined.

For the penetrance, three biological replicates were performed and the mean ± SD for the percentage of animals with the phenotype were calculated and one-way ANOVA followed by Dunnett’s multiple comparisons test (comparison with the wild type) or a Tukey’s honestly significant difference (HSD) test (selected pairwise comparison) was performed using GraphPad Prism 8.

## ACKNOWLEDGEMENTS

We thank Prof. Anton Gartner at the Ulsan National Institute for Science and Technology for providing strains and Prof. Gary YW Chan at HKU for insightful discussion. This work is supported by the National Natural Science Foundation of China (Excellent Young Scientists Fund for Hong Kong and Macau, 32122002), and the Research Grant Council of Hong Kong (GRF 17105523, GRF 17106322, GRF 17113324, and GRF 17107325). Some strains were provided by the Caenorhabditis Genetics Center, which is funded by the National Institutes of Health (NIH) Office of Research Infrastructure Programs (P40 OD010440).

