## Supplementary material for "LEM-3/ANKLE1 nuclease prevents the formation of syncytium between postmitotic sister cells and safeguards neuronal differentiation": Figure S1-S10 and Table S1-S2

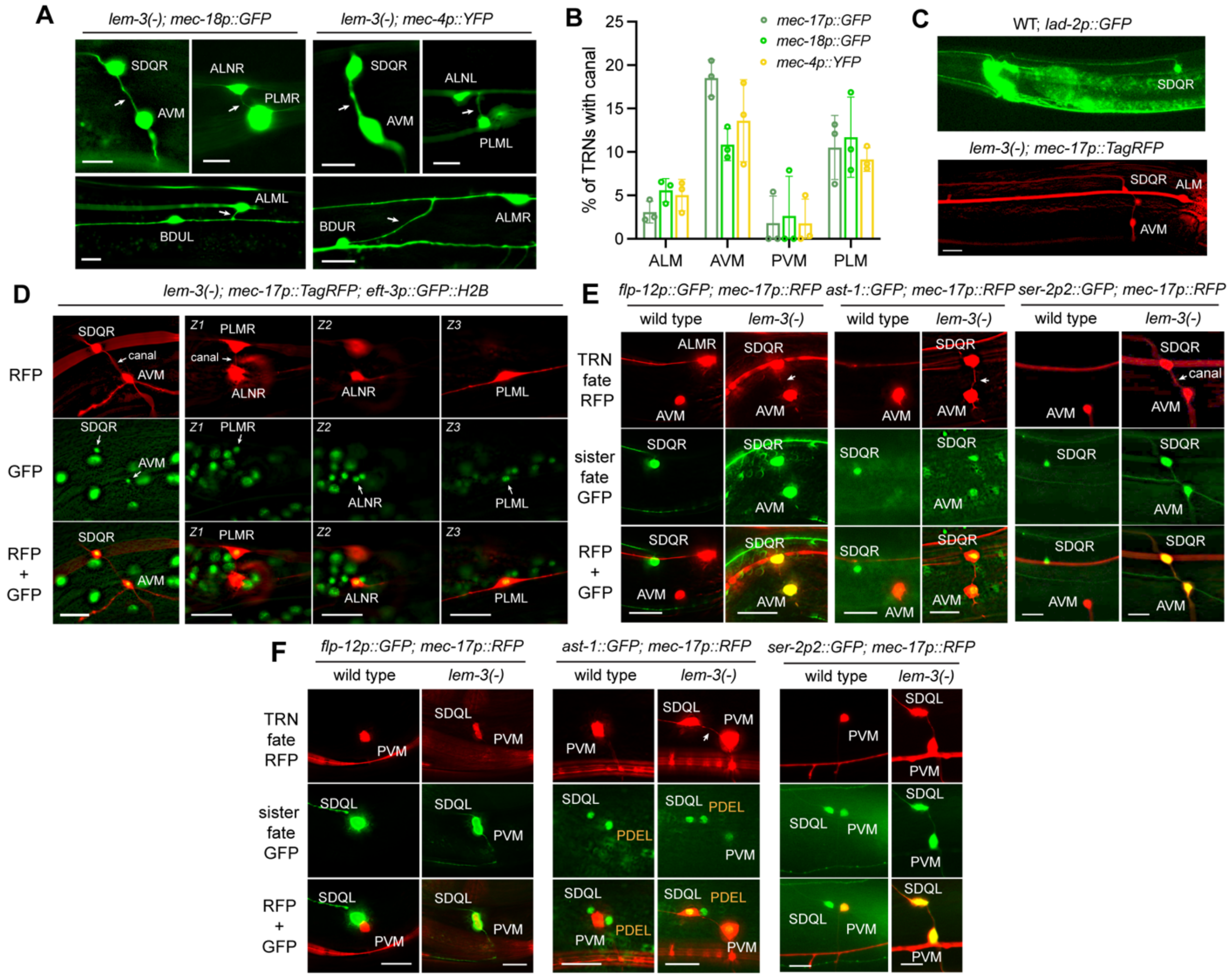


**Figure S1. Binucleate syncytia between TRNs and their sister cells in *lem-3* mutants.** (A) The TRN-sister cell syncytia in *lem-3(unk84)* mutants express TRN markers, *unkEx309[mec-18p::GFP]* and *unkEx313 [mec-4p::YFP]*. Arrows indicate the intercellular canal of the syncytium. Scale bars = 10 µm. (B) The percentage of TRNs showing the syncytium labelled by various TRN markers, including *uIs31[mec-17p::GFP]*, *unkEx309*, and *unkEx313*. (C) Similar morphology of the SDQR neuron in wild-type and *lem-3(-)* mutants that possess the AVM-SDQR syncytium. (D) The two cell bodies in the AVM-SDQR and PLMR-ALNR syncytia both contain histones labelled by *oxTi75[eft-3p::GFP::H2B]*. Arrows point to the histone-containing nuclei. "Z1" and "Z2" indicate two focal planes of the same animal. (E) The AVM-SDQR syncytium expresses various SDQR marker, including *ynIs25[flp-12p::gfp]*, *ast-1::GFP(vlc19)*, and *otIs358[ser-2p2::GFP]*. (F) The PVM-SDQL syncytium is a labelled by SDQL markers, *flp-12p::GFP* and *ast-1::GFP*. Because *ser-2p2p::GFP* is expressed in both PVM and SDQL in the wild-type animals, it cannot be used as a SDQL-specific marker for the syncytium. As expected, the PVM-SDQL syncytium shows *ser-2p2p::GFP* expression.

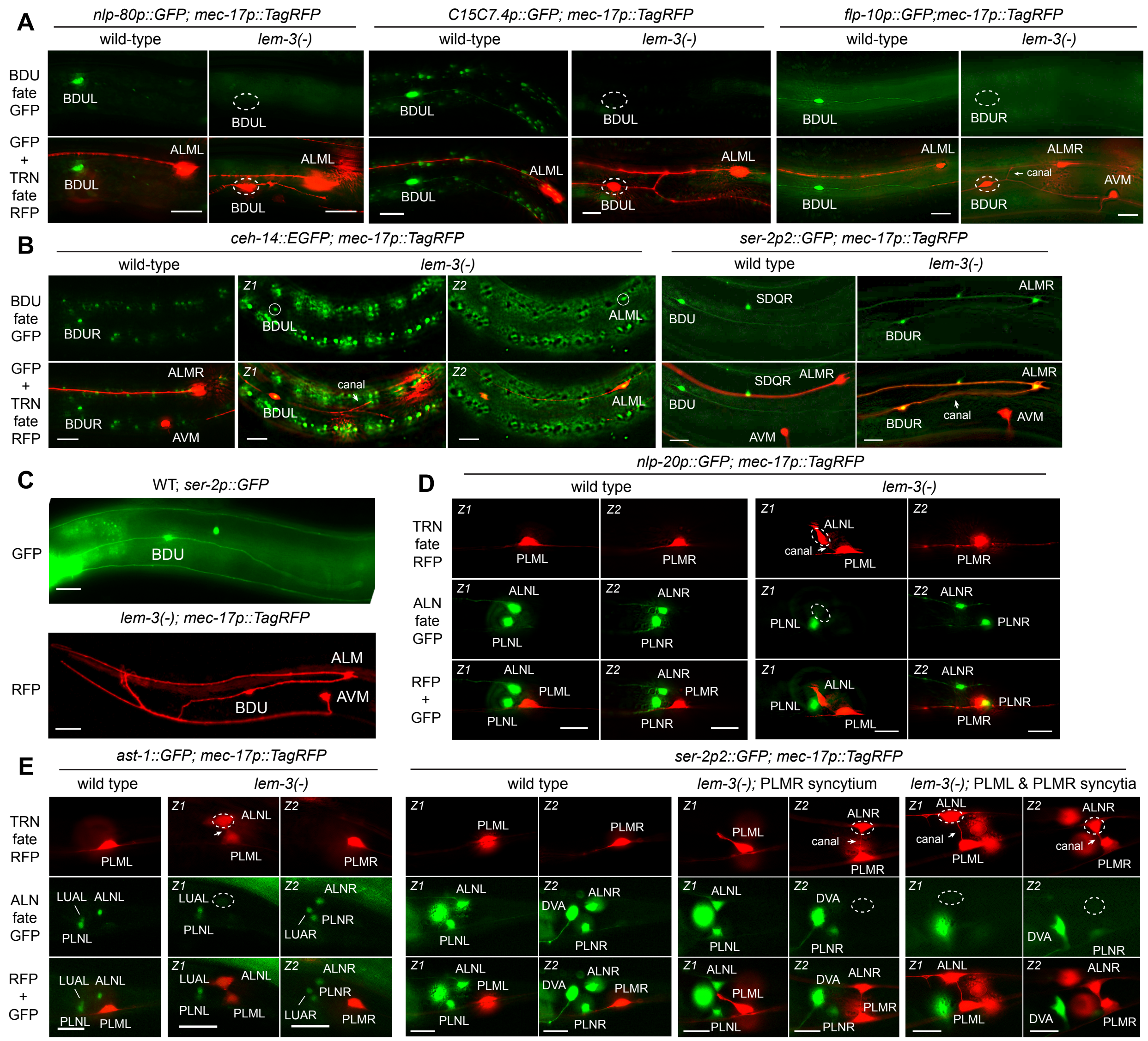


**Figure S2. Repression of sister cell fate markers in ALM-BDU and PLM-ALN syncytia.** (A-B) The lack of expression (dashed circles) of BDU fate markers *unkEx914[nlp-80p::GFP]*, *unkEx917[C15C7.4p::GFP]*, and *otIs92[flp-10::GFP]* in the ALM-BDU syncytium of *lem-3(unk84)* mutants. The persistent expression of BDU fate markers, *wgIs73[ceh-14::EGFP]* (circles) and *otIs358[ser-2p2::GFP]*, in the syncytium. Scale bars = 10 µm. “Z1” and “Z2” indicate two focal planes of the same animal. (C) The comparison of morphology of the BDU neurons labelled by *otIs358* and the BDU of the ALM-BDU syncytium labelled by *uIs115[mec-17p::TagRFP]* in *lem-3(unk84)* mutants. (D-E) The absence of expression (dashed circles) of ALN fate markers *rtEx332[nlp-20p::GFP]*, *ast-1::GFP(vlc19)*, and *otIs358[ser-2p2::GFP]* in the PLM-ALN syncytia. For the case of *otIs358[ser-2p2::GFP]*, two scenarios were shown. One shows only the PLMR-ALNR syncytium and wild-type PLML; the other shows both the PLML-ALNL and the PLMR-ALNR syncytia.

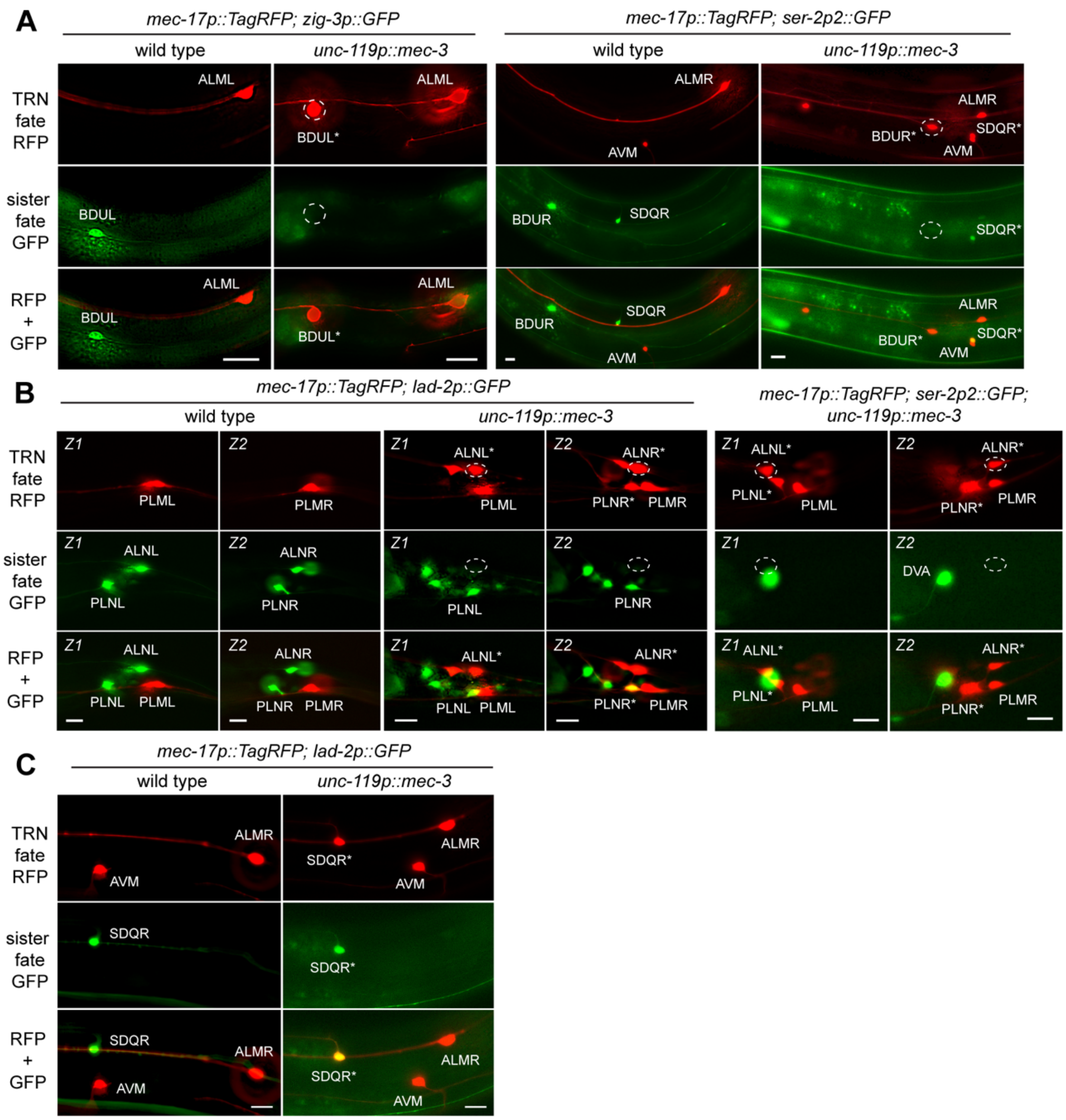


**Figure S3. Misexpression of *mec-3* converts BDU and ALN into TRN-like cells.** (A) BDU neurons express the TRN marker *uIs115* and do not express the BDU fate markers *otIs358[ser-2p2::GFP]* and *otIs14[zig-3p::GFP]* in the animals carrying the *uIs158* transgene. SDQR expresses the TRN marker but also keeps the expression of SDQ marker *otIs358* upon *mec-3* misexpression. (B) ALN neurons express the TRN fate marker *uIs115[mec-17p::TagRFP]* and lose expression of ALN markers *otIs358[ser-2p2::GFP]* and *uIs130[lad-2p::GFP]* in animals carrying the *uIs158[unc-119p::mec-3]* transgene. Wild-type expression pattern of *otIs358[ser-2p2::GFP]* in the tail can be found in Figure S2B for comparison. Dashed circles indicate the lack of expression. Scale bars = 10 μm. Neuron names with an asterisk mean that the neuron abnormally expresses the TRN marker upon forced expression of *mec-3*. (C) SDQR expresses both the TRN marker *uIs115* and the SDQ marker *uIs130[lad-2p::GFP]* in animals carrying the *uIs158* transgene.

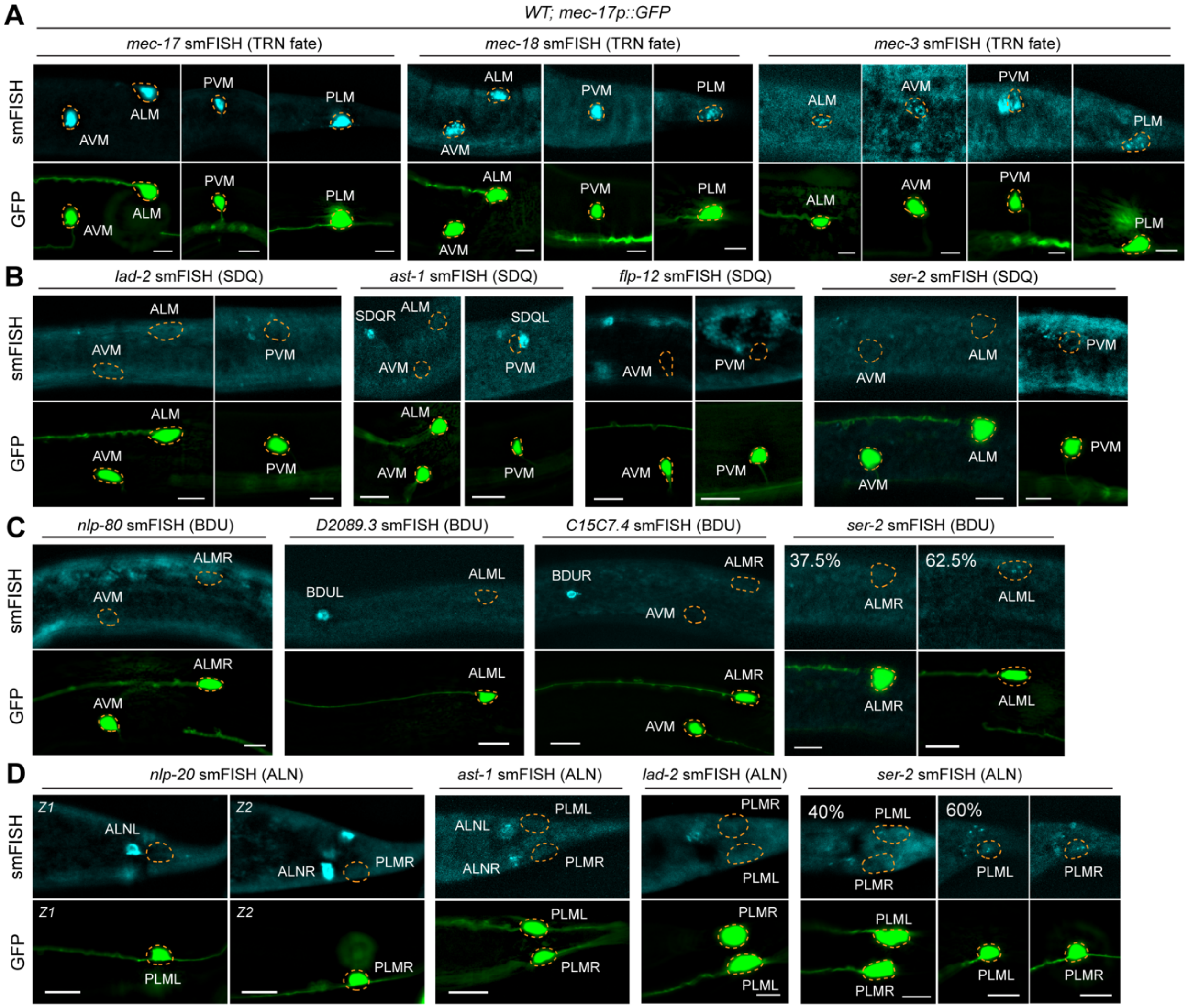


**Figure S4. The lack of endogenous expression of sister cell fate markers in wild-type TRNs.** (A) smFISH signals for the mRNAs of TRN genes *mec-17*, *mec-18*, and *mec-3* in wild-type animals. Dashed orange circles indicate the cell bodies. The *uIs31[mec-17p::GFP]* transgene was used to label the TRNs. Scale bars = 5 µm. (B) Lack of smFISH signals for SDQ marker genes *lad-2*, *ast-1*, and *flp-12* in AVM and PVM neurons. SDQR marker *ser-2* mRNAs are not found in AVM but are found in 100% (*n* = 10) of PVM neurons. (C) smFISH signals for BDU fate markers *nlp-80*, *D2089.3*, and *C15C7.4* are not observed in ALM neurons. *ser-2* smFISH signals are found in 62.5% (*n* = 40) of ALM neurons. (D) smFISH signals for ALN fate markers *nlp-20*, *ast-1*, and *lad-2* are not found in PLM neurons. *ser-2* smFISH signals are found in 60% (*n* = 40) of PLM neurons.

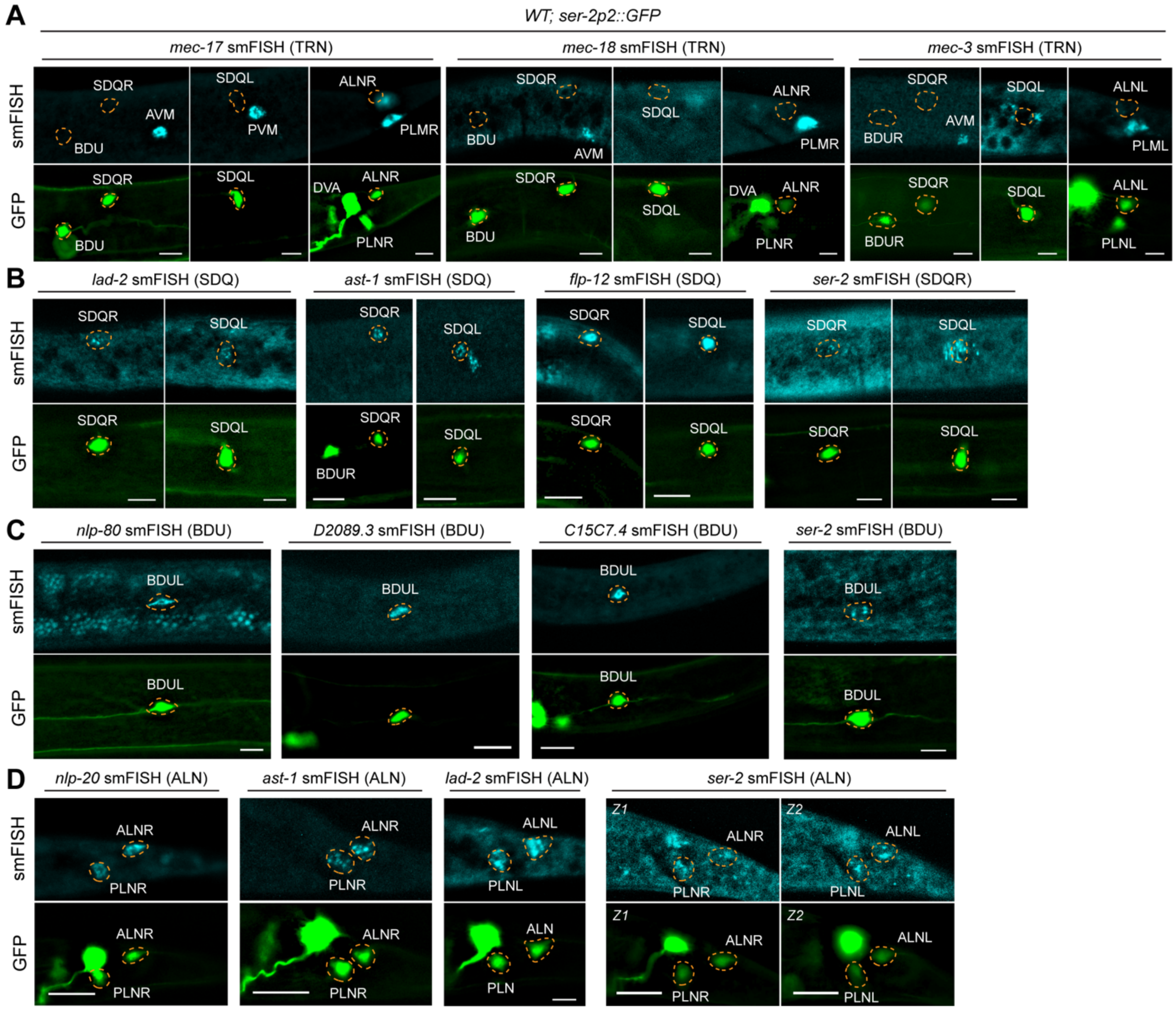


**Figure S5. Confirmation of the mRNA expression of TRN sister cell fate markers in wild-type animals.** (A) smFISH signals for the TRN fate markers *mec-17*, *mec-18*, and *mec-3* are not found in the sister cells of TRNs in the wild-type animals. Dashed orange circles indicate the cell bodies. The *otIs358[ser-2p2::GFP]* transgene was used to label TRN sister cells. Scale bars = 5 µm. (B) smFISH signals for the SDQ marker genes *lad-2*, *ast-1*, and *flp-12* in SDQL/R neurons. *ser-2* only serves as a SDQR marker, although its mRNAs are found in both SDQL and SDQR. (C) smFISH signals for BDU fate markers *nlp-80*, *D2089.3*, *C15C7.4*, and *ser-2* in BDUL/R neurons. (D) smFISH signals for ALN fate markers *nlp-20*, *ast-1*, *lad-2*, and *ser-2* in ALNL/R neurons.

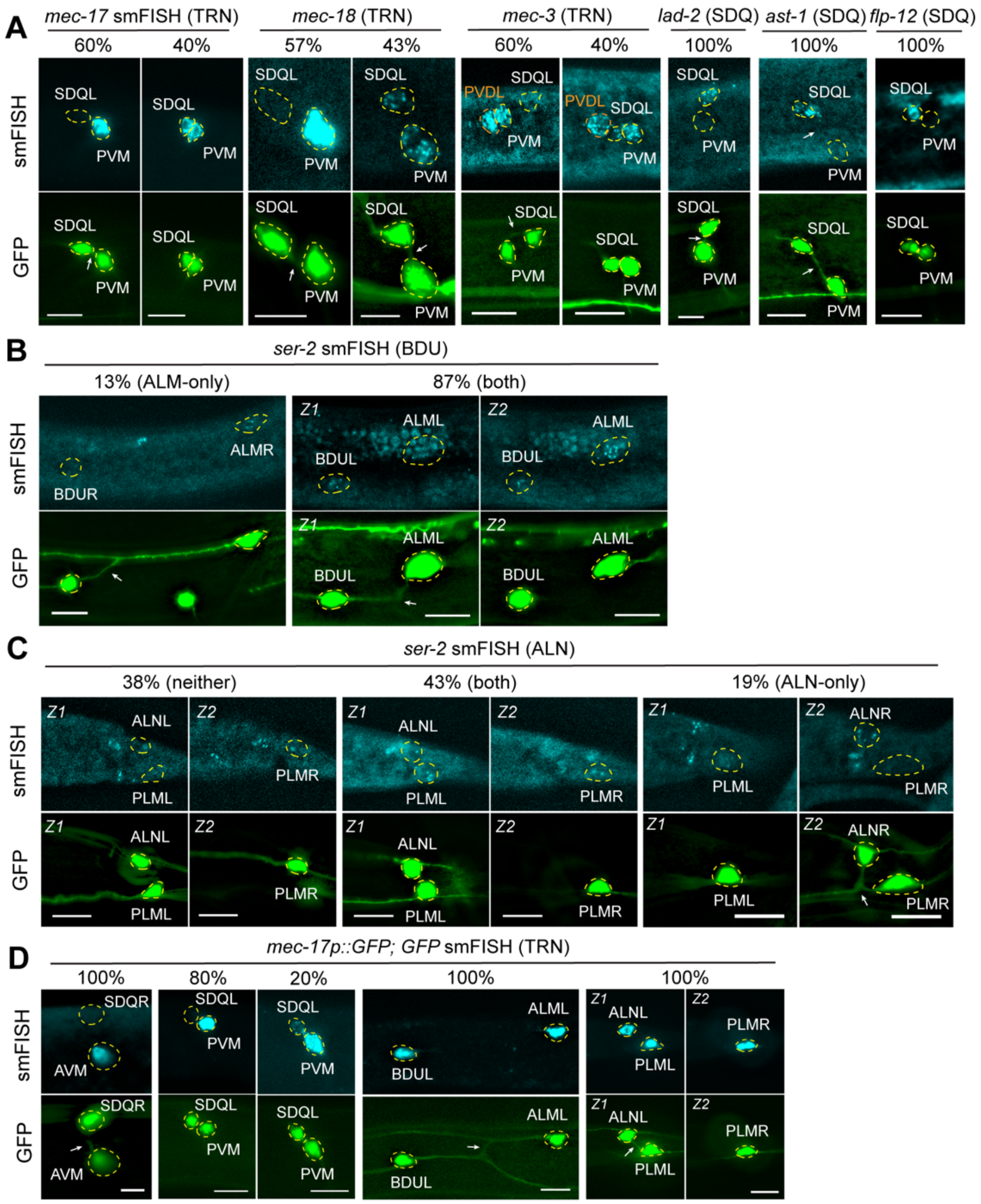


**Figure S6. Additional evidence for the activation of endogenous fate markers in the two cell bodies of the syncytia.** (A) smFISH signals for the mRNAs of TRN genes (*mec-17*, *mec-18*, and *mec-3*) and SDQL genes (*lad-2*, *ast-1*, and *flp-12*) in the two cell bodies of the PVM-SDQL syncytia in *lem-3(unk84)* mutants ($n \geq 10$). Dashed circles indicate the cell bodies. "Z1" and "Z2" indicate different focal planes of the same animal. Arrows point to the intercellular canal. Scale bars = 5 µm. (B) *ser-2* smFISH signals are absent in the BDU cell body for 13% of the ALM-BDU syncytia ($n = 15$). (C) *ser-2* smFISH signals are absent in the ALN cell body for 38% of the PLM-ALN syncytia ($n = 21$). (D) smFISH signals for GFP mRNAs in the *lem-3(unk84); uIs31[mec-17p::GFP]* strain. Percentage of cells having the shown smFISH staining patterns ($n \geq 10$).

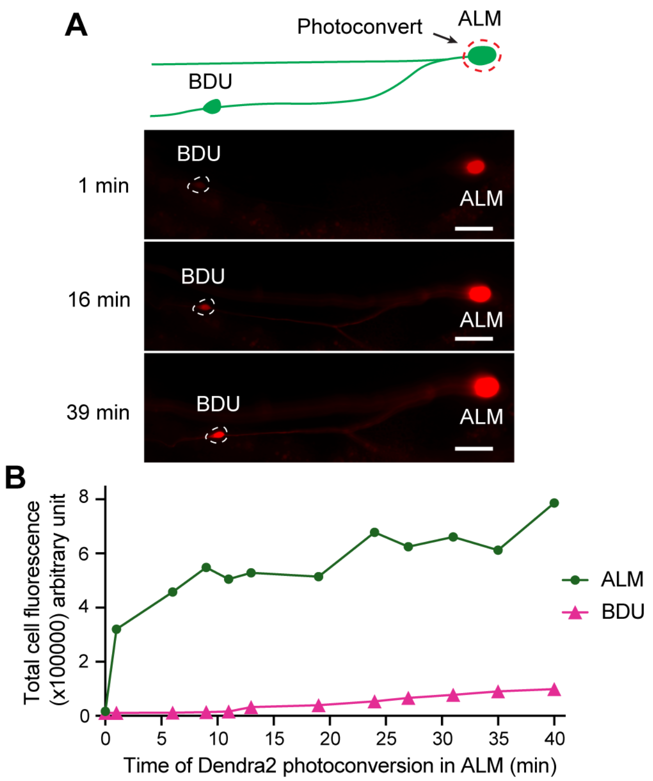


**Figure S7. Diffusion of proteins between the two cell bodies of the ALM-BDU syncytium.** (A) Photoconversion of dendra2 in the ALM cell body of the ALM-BDU syncytium led to the increase of red fluorescence in BDU neuron (dashed circle). *lem-3(unk84); unkEx271[mec-17p::dendra2]* was used in this experiment. Scale bars = 10 µm. (B) Quantification of the red fluorescence in both ALM and BDU cell bodies throughout the course of the photoconversion experiment.

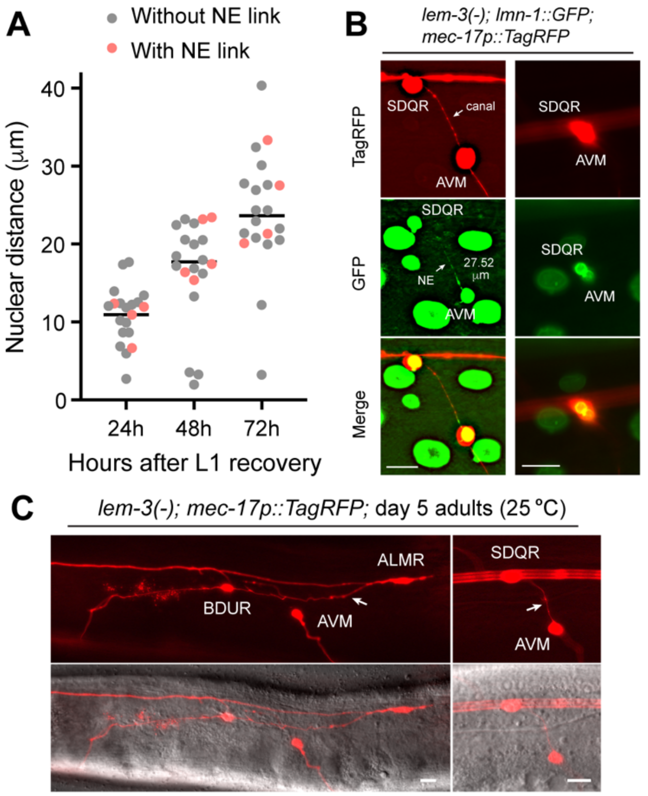


**Figure S8. Intercellular canal and nuclear envelope connection in the TRN-sister cell syncytia are maintained throughout development and aging.** (A) Quantification of the distance between the two nuclei of the AVM-SDQR syncytia at different developmental stages. *lem-3(unk84); uIs115[mec-17p::RFP]; ccIs4810[lmn-1p::lmn-1::GFP]* was used to label both the cytosol and nuclear envelope. The syncytia with nuclear envelope (NE) link were marked in red in the plot. (B) Representative images of syncytia with a long NE link or with unseparated nuclei at 72 hours post L1 arrest recovery are shown. Scale bars = 10 μm. (C) Intercellular canal (arrows) of ALM-BDU and AVM-SDQR syncytia at the day 5 adult stage in *lem-3(unk84)* mutants.

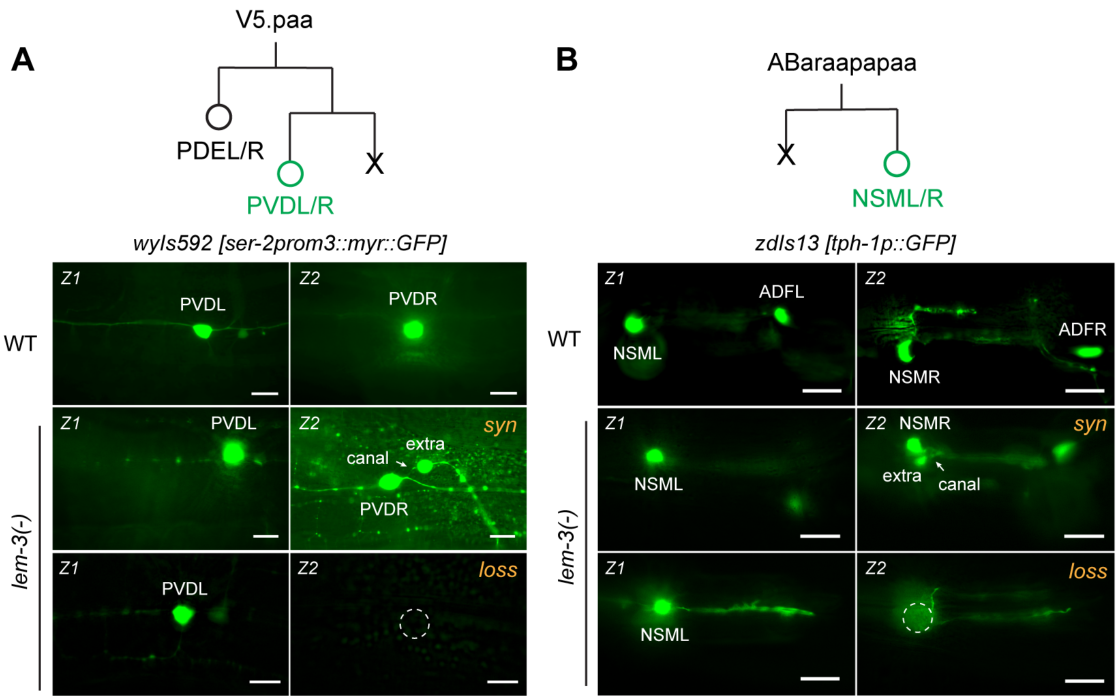


**Figure S9. Formation of syncytia between neurons and their apoptotic sisters.** (A) The sister cell of PVDL/R is apoptotic. In the *lem-3(unk84); wyIs592[ser-2p3::myr::GFP]* animals, which labels the PVD neuron, both syncytia of PVD with an extra cell ("*syn*") and the loss of PVD labelling ("*loss*") are found. For the former phenotype, arrows point to the canal and for the latter phenotype, dashed circle indicates the absence of GFP signal. Scale bars = 5 µm. "Z1" and "Z2" indicate different focal planes of the same animal. (B) The syncytia and the loss of fate marker phenotypes for NSM neurons in the *lem-3(-)* mutants.

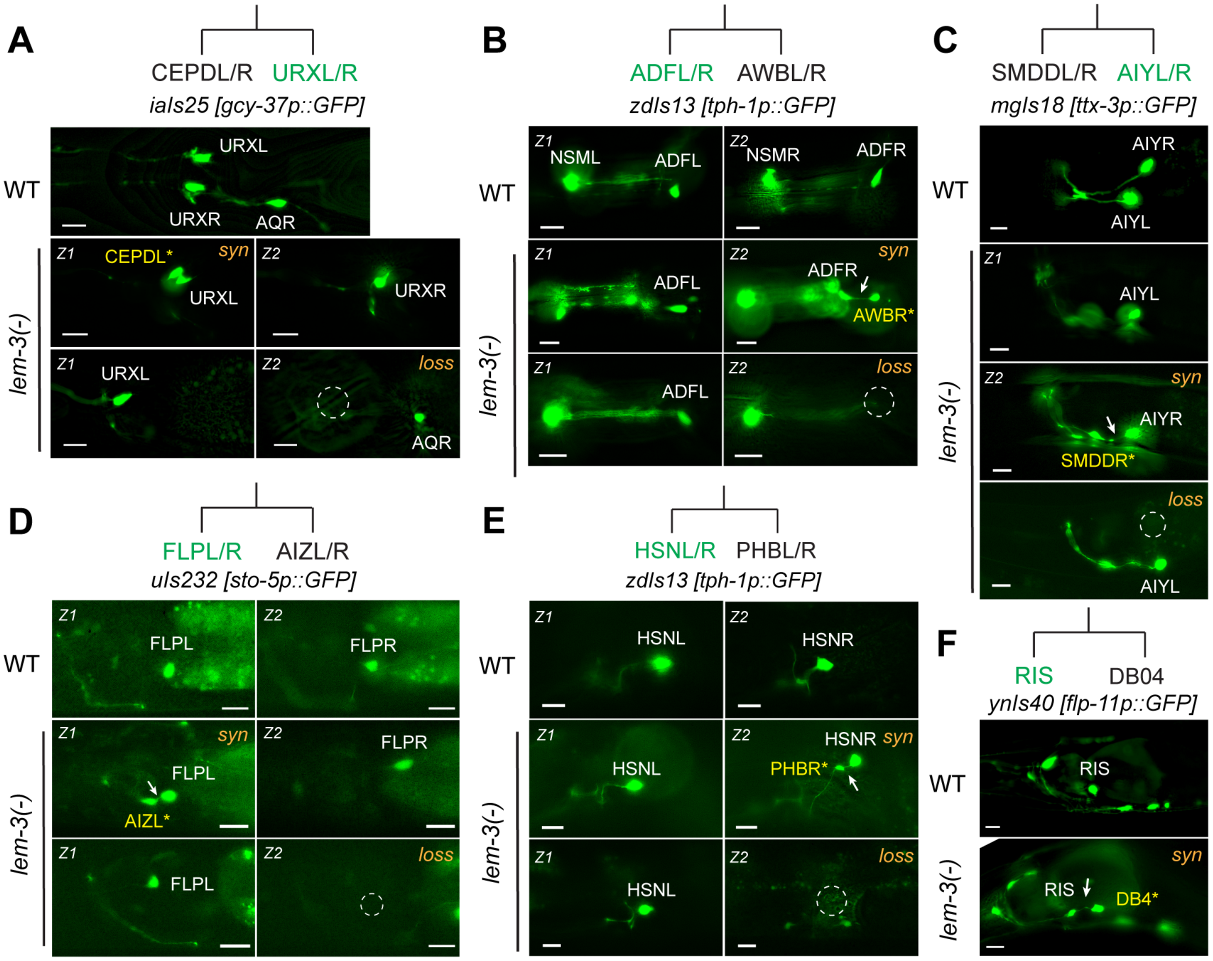


**Figure S10. Formation of syncytia between multiple sister cell pairs in *lem-3(-)* mutants.** (A-E) Representative images of syncytia ("*syn*") and fate marker loss ("*loss*" and dashed circles) phenotypes for URXL/R, ADFL/R, AIYL/R, FLPL/R, and HSNL/R neurons in *lem-3(unk84)* mutants. In syncytia, sister cells abnormally expressing markers are labelled with yellow names; arrows indicate the intercellular canal. Quantification of penetrance is shown in Figure 7F. Scale bars = 5 µm. "Z1" and "Z2" mean two focal planes of the same animal. (F) The syncytium phenotype for the RIS neuron in *lem-3(unk84)* mutants treated with 10 µM ICRF-193.

**Table S1. Strains, constructs, and primers used in this study.**

| Strain list | |
|---|---|
| **Strain name** | **Genotype** |
| CGZ1053 | *lem-3(unk84) I; oxTi75 [eft-3p::GFP::H2B::tbb-2 3'UTR + unc-18(+)] II; uIs115 [mec-17p::RFP] IV* |
| CGZ1054 | *lem-3(unk84) I; uIs115 [mec-17p::RFP] IV; unkEx242 [lem-3p::lem-3::lem-3 3'UTR (9kb) + ceh-22p::GFP]* |
| CGZ1158 | *uIs115 [mec-17p::RFP] IV; uIs130 [lad-2p::GFP] X; uIs158 [unc-119p::mec-3]* |
| CGZ1159 | *lem-3(unk84) I; ayIs9 [egl-17p::GFP + dpy-20(+)] II; casIs131 [egl-17p::mCherry::zen-4]* |
| CGZ1160 | *lem-3(unk14) I; unkEx271 [mec-17p::dendra2::unc-54 3'UTR + ceh-22p::GFP]* |
| CGZ1198 | *lem-3(unk84) I; uIs115 [mec-17p::RFP] IV; rtEx332 [nlp-20p::GFP + lin-15(+)]* |
| CGZ1223 | *lem-3(unk84) I; wzIs112 [gcy-9p::GFP]* |
| CGZ1312 | *lem-3(unk84) I; unkEx309 [mec-18p::GFP + myo-3p::mCherry]* |
| CGZ1326 | *lem-3(unk84) I; uIs115 [mec-17p::RFP] IV; unkEx313 [mec-4p::YFP + myo-3p::mCherry]* |
| CGZ1383 | *lem-3(unk84) I; uIs115 [mec-17p::RFP] IV; ccIs4810 [(pJKL380.4) lmn-1p::lmn-1::GFP::lmn-1 3'UTR + (pMH86) dpy-20(+)] X* |
| CGZ1384 | *lem-3(unk84) I; uIs115 [mec-17p::RFP] IV; uIs130 [lad-2p::GFP] X* |
| CGZ1585 | *lem-3(unk84) I; unc-119(ed3) III; uIs115 [mec-17p::RFP] IV; unkEx170 [maph-1.1p::maph-1.1::GFP::maph-1.1 3'UTR_2 + unc-119(+)]* |
| CGZ1656 | *lem-3(unk84) I; uIs115[mec-17p::RFP] IV; wgIs55 [mec-3::TY1::EGFP::3xFLAG(92C12) + unc-119(+)]* |
| CGZ1744 | *lem-3(unk84) I; ynIs78 [flp-8p::GFP] X* |
| CGZ1745 | *lem-3(unk84) I; ynIs40 [flp-11p::GFP] V* |
| CGZ1749 | *lem-3(unk84) I; oxIs12 [unc-47p::GFP + lin-15(+)] X* |
| CGZ1750 | *lem-3(unk84) I; zdIs13 [tph-1p::GFP] IV* |
| CGZ1751 | *lem-3(unk84) I; oyIs18 [gcy-8::GFP] X* |
| CGZ1828 | *lem-3(unk84) I; unc-119(ed3) III; uIs115 [mec-17p::RFP] IV; unkEx573 [mec-18p::MS2BP::GFP::2xGGS::NLS + unc-119(+)]* |
| CGZ1832 | *lem-3(unk84) I; uIs31 [mec-17p::GFP] III; uIs165 [mec-4::TagRFP; inx-20p::GFP]* |
| CGZ1858 | *lem-3(unk84) I; uIs115 [mec-17p::RFP] IV; jsIs821 [mec-7p::GFP::rab-3] X* |
| CGZ1859 | *lem-3(unk84) I; uIs115 [mec-17p::RFP] IV; jsIs609 [mec-7p::mitoGFP + lin-15(+)] X* |
| CGZ1935 | *lem-3(unk84) I; uIs31 [mec-17p::GFP] III; bqSi226 [emr-1p::emr-1::mCherry + unc-119(+)] IV* |
| CGZ2001 | *lem-3(unk84) I; unc-119(ed3) III; uIs115 [mec-17p::RFP] IV; unkEx631 [mec-17p::PLCdPH::GFP + unc-119(+)]* |
| CGZ2046 | *lem-3(unk84) I; unc-119(ed3) III; uIs115 [mec-17p::RFP] IV; unkEx636 [mec-17p::LiveAct::GFP::mec-17 3'UTR + unc-119(+)]* |
| CGZ2057 | *lem-3(unk84) I; unc-119(ed3) III; uIs115 [mec-17p::RFP] IV; unkEx642 [mec-18p::mec-6::GFP::unc-54 3'UTR + unc-119(+)]* |
| CGZ2144 | *lem-3(unk84) I; vtIs1 [dat-1p::GFP + rol-6(su1006)] V* |

| | |
|---|---|
| CGZ2213 | *lem-3(unk84) I; uIs232 [sto-5p::GFP]* |
| CGZ2258 | *otIs358 [ser-2p2::GFP + pha-1(+)] IV; uIs134 [mec-17p::RFP] V* |
| CGZ2264 | *lem-3(unk84) I; uIs115 [mec-17p::RFP] IV; otIs92 [flp-10p::GFP]* |
| CGZ2286 | *otIs358 [ser-2p2::GFP + pha-1(+)] IV; uIs134 [mec-17p::RFP] V; uIs158 [unc-119p::mec-3]* |
| CGZ2330 | *lem-3(unk84) I; uIs115 [mec-17p::RFP] IV; uIs72 [pCFJ90(myo-2p::mCherry) + unc-119p::sid-1 + mec-18p::mec-18::GFP]* |
| CGZ2468 | *lem-3(unk84) I; uIs115 [mec-17p::RFP] IV; otIs14 [zig-3p::GFP + rol-6(su1006)]* |
| CGZ2522 | *uIs115 [mec-17p::RFP] IV; otIs14 [zig-3p::GFP + rol-6(su1006)]* |
| CGZ2612 | *uIs115 [mec-17p::RFP] IV; uIs158 [unc-119p::mec-3]; otIs14 [zig-3p::GFP + rol-6(su1006)]* |
| CGZ2613 | *uIs115 [mec-17p::RFP] IV; wgIs73 [ceh-14::TY1::EGFP::3xFLAG + unc-119(+)]* |
| CGZ2614 | *lem-3(unk84) I; uIs115 [mec-17p::RFP] IV; wgIs73 [ceh-14::TY1::EGFP::3xFLAG + unc-119(+)]* |
| CGZ2677 | *lem-3(unk84) I; uIs115 [mec-17p::RFP] IV; ynIs25 [flp-12p::GFP]* |
| CGZ2678 | *uIs115 [mec-17p::RFP] IV; ynIs25 [flp-12p::GFP]* |
| CGZ2679 | *lem-3(unk84) I; otIs181 [dat-1p::mCherry + ttx-3p::mCherry] III; iaIs25 [gcy-37p::GFP + unc-119(+)]* |
| CGZ2680 | *otIs181 [dat-1p::mCherry + ttx-3p::mCherry] III; iaIs25 [gcy-37p::GFP + unc-119(+)]* |
| CGZ2681 | *wyIs592 [ser-2p3::myr::GFP + odr-1p::RFP] III; ced-3(n717) IV* |
| CGZ2686 | *lem-3(unk84) I; unc-119(ed3) III; uIs115 [mec-17p::RFP] IV; unkEx914 [nlp-80p::GFP::unc-54 3'UTR + unc-119(+)]* |
| CGZ2692 | *lem-3(unk84) I; unc-119(ed3) III; uIs115 [mec-17p::RFP] IV; unkEx917 [C15C7.4p::GFP::unc-54 3'UTR + unc-119(+)]* |
| CGZ2760 | *ast-1(vlc19 [ast-1::GFP]) II; uIs115 [mec-17p::RFP] IV* |
| CGZ2761 | *lem-3(unk84) I; ast-1(vlc19 [ast-1::GFP]) II; uIs115 [mec-17p::RFP] IV* |
| CGZ28 | *lem-3(unk14) I; uIs134 [mec-17p::RFP] V* |
| CGZ2844 | *uIs115 [mec-17p::RFP] IV; rtEx332 [nlp-20p::GFP + lin-15(+)]* |
| CGZ2845 | *uIs115 [mec-17p::RFP] IV; unkEx917 [C15C7.4p::GFP::unc-54 3'UTR + unc-119(+)]* |
| CGZ2846 | *uIs115 [mec-17p::RFP] IV; unkEx914 [nlp-80p::GFP::unc-54 3'UTR + unc-119(+)]* |
| CGZ2847 | *uIs115 [mec-17p::RFP] IV; otIs92 [flp-10p::GFP]* |
| CGZ32 | *lem-3(mn155) I; uIs134 [mec-17p::RFP] V* |
| CGZ33 | *lem-3(unk14) I; uIs31 [mec-17p::GFP] III* |
| CGZ599 | *lem-3(unk84) I; uIs115 [mec-17p::RFP] IV* |
| CGZ601 | *lem-3(unk85) I; uIs115 [mec-17p::RFP] IV* |
| CGZ667 | *zdIs13 [tph-1p::GFP] IV* |
| CGZ742 | *lem-3(unk84) I; iaIs25 [gcy-37p::GFP + unc-119(+)]* |
| CGZ788 | *lem-3(unk84) I; wyIs592 [ser-2p3::myr::GFP + odr-1p::RFP] III* |
| CGZ790 | *lem-3(unk84) I; mgIs18 [ttx-3p::GFP] IV* |

| | | |
|---|---|---|
| CGZ791 | *lem-3(unk84) I; ntIs1 [gcy-5p::GFP + lin-15(+)] V* | |
| CGZ809 | *lem-3(unk84) I; otIs358 [ser-2p2::GFP + pha-1(+)] IV; uIs134 [mec-17p::RFP] V* | |
| CGZ810 | *air-2(or207) I; uIs115 [mec-17p::RFP] IV* | |
| CGZ830 | *lem-3(gt3310 [eGFP::STag::lem-3 [S192A S194A]]) I; uIs115 [mec-17p::RFP] IV* | |
| CGZ831 | *lem-3(gt3311 [eGFP::STag::lem-3 [Y556A G558A]]) I; uIs115 [mec-17p::RFP] IV* | |
| CGZ866 | *lem-3(unk84) I; otIs92 [flp-10p::GFP]* | |
| EG1285 | *lin-15B&lin-15A(n765); oxIs12 [unc-47p::GFP + lin-15(+)] X* | |
| FQ383 | *wzIs112 [gcy-9p::GFP]* | |
| NY2040 | *ynIs40 [flp-11p::GFP] V* | |
| NY2078 | *ynIs78 [flp-8p::GFP]* | |
| OH10765 | *pha-1(e2123) III; otIs358 [ser-2p2::GFP + pha-1(+)]* | |
| OH3192 | *ntIs1 [gcy-5p::GFP + lin-15(+)] V* | |
| OH4841 | *otIs92 [flp-10p::GFP]* | |
| OH99 | *mgIs18 [ttx-3p::GFP] IV* | |
| PY1322 | *oyIs18 [gcy-8::GFP] X* | |
| TG2435 | *vtIs1 [dat-1p::GFP + rol-6(su1006)] V* | |
| TU4065 | *uIs115 [mec-17p::RFP] IV* | |
| TU4084 | *uIs115 [mec-17p::RFP] IV; uIs130 [lad-2p::GFP] X* | |
| TU4864 | *uIs232 [sto-5p::GFP]* | |
| TU6565 | *uIs31 [mec-17p::GFP] III* | |
| TV15911 | *wyIs592 [ser-2p3::myr::GFP + odr-1p::RFP] III* | |
| ZG610 | *iaIs25 [gcy-37p::GFP + unc-119(+)]* | |
| **Construct list** | | |
| **Construct name** | **Construct details** | **Comments** |
| CGZ#1232 | *nlp-80p::GFP::unc-54 3'UTR* | Used to create CGZ2686 |
| CGZ#1261 | *C15C7.4p::GFP::unc-54 3'UTR* | Used to create CGZ2692 |
| CGZ#276 | *maph-1.1p::maph-1.1::GFP::maph-1.1 3'UTR_2* | Used to create CGZ1585 |
| CGZ#412 | *mec-17p::dendra2::unc-54 3'UTR* | Used to create CGZ1160 |
| CGZ#598 | *mec-18p::MS2BP::GFP::2xGGS::NLS* | Used to create CGZ1828 |
| CGZ#79 | *lem-3p::lem-3::lem-3 3'UTR* | Used to create CGZ1054 |
| CGZ#801 | *mec-18p::mec-6::GFP::unc-54 3'UTR* | Used to create CGZ2057 |
| CGZ#810 | *mec-17p::LiveAct::GFP::mec-17 3'UTR* | Used to create CGZ2046 |
| TU#1739 | *mec-4p::YFP* | Used to create CGZ1326 |
| TU#2041 | *mec-17p::PLCdPH::GFP* | Used to create CGZ2001 |
| TU#924 | *mec-18p::GFP::unc-54 3'UTR* | Used to create CGZ1312 |
| **Primer list** | | |
| **Oligo Name** | **Sequence (5'-3')** | **Comments** |

| | | |
|---|---|---|
| nlp-80p_f | GATACGCTAACAACTTGGAACCCAACTGC GATACTCCAAC | Used to create CGZ#1232 |
| nlp-80p_r | AGTTCTTCTCCTTTACTCATTCCTGAAAAA ATGATTGAAATATGAGAAGAAACAG | Used to create CGZ#1232 |
| C15C7.4p_f2 | GATACGCTAACAACTTGGAATAGTACGCTC CAGTTAGGAAAGACG | Used to create CGZ#1261 |
| C15C7.4p_r | AGTTCTTCTCCTTTACTCATTCTGCAGAAA AGTGGCATTAGTAAATTTG | Used to create CGZ#1261 |
| GFP-HA-f | AGTAAAGGAGAAGAACTTTTCACTGG | Used to create CGZ#276 |
| maph-1.1-KLD-R | GAGCAAATCGACTCTGGCCA | Used to create CGZ#276 |
| mec17p_dendra2_f | AGTTGTGAGACGATTCGATCATGAATACCC CTGGAATTAACCTTATTAAGG | Used to create CGZ#412 |
| dendra2_unc54_r | TTGGAATTCTACGAATGCTACCACACTTGA CTTGGTAGAGGACT | Used to create CGZ#412 |
| NLS_GFP_f1 | CCAAAGAAGAAACGCAAAGTATAGCATTC TAGAATTCCACTGAGCG | Used to create CGZ#598 |
| GFP_2xGGS_r1 | GGAGCCACCTGAGCCTCCTTTGTATAGTTC ATCCATGCCATGTGTAATCC | Used to create CGZ#598 |
| pUC57_lem-3_9kb_f | GTTGTAAAACGACGGCCAGTCGTTGGCAC TGAGCGTTATCG | Used to create CGZ#79 |
| pUC57_lem-3_r | CAGGAAACAGCTATGACCATAGCTGTTGG TGGAGATGTGTTC | Used to create CGZ#79 |
| mec-18p_mec-6_f | CTTCCAAGAAGGTTGTGAGCATGGGTCTC AATCGGCAGC | Used to create CGZ#801 |
| mec-6_GFP_r | TTGGCCAATCCCGGGGATCCTGTAATATAG AATGCGTAAGATCACAATGAAG | Used to create CGZ#801 |
| LActin_GGS_f | CAAGAAGTTCGAATCTATCTCCAAGGAGG AGGGCGGCAGTGGAGGATCT | Used to create CGZ#810 |
| mec-17p_LActin_r | ATAAGATCAGCGACTCCCATGATCGAATCG TCTCACAACTGATCCA | Used to create CGZ#810 |

**Table S2. smFISH and smiFISH probe sets used in this study.**

| ***mec-17* smFISH probe set** | |
|---|---|
| Probe # | Probe Sequence (5' to 3') |
| 1 | gtctcacaactgatccatcg |
| 2 | cgtcgacttgcatgatcgaa |
| 3 | gagttgaggaccgagaatcg |
| 4 | ctctcataggatccaaacga |
| 5 | caattggatcctgcaattgc |
| 6 | gcaagattatcgattgcttc |
| 7 | aaggcaatgagctgagagtt |
| 8 | ttgtcaaaggagtacgaagc |
| 9 | cggagttgataagcttctca |
| 10 | ctcttcttcatcatatttcc |
| 11 | gaatcccatgagtcgactaa |
| 12 | acttctttcttcctactttc |
| 13 | agtctgcatttgagagtcat |
| 14 | gaagacacagaatctctccc |
| 15 | ggcgttgacatgagaagtga |
| 16 | agaatttgttgtccaactcc |
| 17 | gtgttcctgagagaacatgt |
| 18 | gagccaattgataaggttca |
| 19 | gtccgtacttttgtgacata |
| 20 | ttggtattctgccatacagg |
| 21 | gagctcctcgaaaacaacga |
| 22 | ttttcagccgacaaagctag |
| 23 | tggtggaggtttttcgattc |
| 24 | aatcgacgtggagtcattgg |
| 25 | ggtatcagtcattccagttc |
| 26 | taacagcgtgttgaagccaa |
| 27 | atttcccttgctctgatgac |
| 28 | ttgtggtgtcatgtcagcat |
| 29 | gagctctattgctcaaagca |
| 30 | tgtgagcttttcttgccttg |
| 31 | cacaatggcttgctcgacag |
| 32 | tcaatgtccacatccgaatc |
| 33 | gcgagattaaacagatggca |
| 34 | gtaagtgtggtggatgttca |
| 35 | ttatttgagggtgttgatga |
| ***mec-18* smFISH probe set** | |
| Probe # | Probe Sequence (5' to 3') |
| 1 | ctcacaaccttcttggaagg |
| 2 | cagttttgtcccaacttatt |

| | |
|---|---|
| 3 | attgccaaattactcccatg |
| 4 | atatctccattgttttccac |
| 5 | tcacaccagtttcgtatttc |
| 6 | agctcttttaatcttgtcgc |
| 7 | actctacttccagacgttac |
| 8 | aaatatcgcttgtccggttg |
| 9 | gtttacagcgactgcagaac |
| 10 | ggtcaaccaacgtccaaatt |
| 11 | atctgagcctttcttctaac |
| 12 | aacatgcttgatacgtcctc |
| 13 | tcaggacgtcttcaatttgc |
| 14 | gtgcactgtccacattaatt |
| 15 | ggtggtttttgtttcacatc |
| 16 | gcactggggttagatttatc |
| 17 | tgacacgggcgatgtcaaat |
| 18 | ctttggattggccagaaact |
| 19 | ctctctacgtctccattaag |
| 20 | gcacttcttcactatctaca |
| 21 | ttttgccctgaaacttgttg |
| 22 | aacgatggctaagttcagct |
| 23 | agcgttcttttgtttgaacc |
| 24 | tgataagtgcgtagtaggca |
| 25 | caatcgattcgactgcttgg |
| 26 | gcatcacgtttatctggtta |
| 27 | atccagtgtacaatagctgg |
| 28 | ttcattgcagcagctgaatt |
| 29 | atcccgaatgattgtcggaa |
| 30 | ggatttcccacagattcaat |
| 31 | tttgaacagcattcctggta |
| 32 | tccgagcacaatgatctgac |
| 33 | agttcagacgtagcttttgg |
| 34 | acttcgtcataaaacccagc |
| 35 | gtccttgattcgatctagta |
| 36 | gggcaaatcatagttccttt |
| 37 | atcatctattccagcatgtg |
| 38 | atgatcctgtcttccaacaa |
| 39 | aaacgctgcaggaacttctc |
| 40 | aaagtggatgctgagcgttc |
| 41 | tttccagaaacatactgccg |
| 42 | cacgcagctctttgaaagtt |
| 43 | ccacatacacttctgggaat |
| 44 | tgtcgaaggtttcgtctcaa |
| 45 | ccactttggaatttgttcgt |

| | |
|---|---|
| 46 | tttttgctccatctttgttg |
| 47 | cgagccgttaactttctttg |
| 48 | ccaattcacaagcaggagtt |
| ***mec-3* smFISH probe set** | |
| Probe # | Probe Sequence (5' to 3') |
| 1 | ctgtaacaaatcacccgtca |
| 2 | aatgcgcgaaattgtggcta |
| 3 | gctttgactctaacatttcc |
| 4 | accatactaatcgcactgag |
| 5 | gattttcatgatcaactcca |
| 6 | tgctcattgcagcaattaca |
| 7 | tacgatcgattgtccattcg |
| 8 | ggagactcacatattgtaca |
| 9 | tccaaaaacacttttcggcc |
| 10 | gactacaataaattctccca |
| 11 | gcacaccgatgaatactgtg |
| 12 | tgacactcctttcttgcaac |
| 13 | gccttcagtttatataccat |
| 14 | acattccacatggaacacga |
| 15 | gataaatgccgtccacacaa |
| 16 | tcaacaagaatttgttcccc |
| 17 | caggatacggtcatcattgt |
| 18 | gttgtcatccatttgaggag |
| 19 | tcaactgcagatcctattgc |
| 20 | tttctgtcgaacaagatggt |
| 21 | catcaattggatatggggca |
| 22 | gtcaacctcctttttaactt |
| 23 | atgctcaaagttgtatccat |
| 24 | tcatcatcacagaaatcgga |
| 25 | gcgtctttttagcattcgtg |
| 26 | gtttgattgtcgttctaggt |
| 27 | tttcattcagcacgtccaac |
| 28 | ggctttggtgtattcgagaa |
| 29 | atttggctcgagcatgtttc |
| 30 | cttaaaccggtctccaatgc |
| 31 | ccaaacctgaattactctca |
| 32 | tccttacttctccggttttg |
| 33 | tgcacaaatgtttgagcctc |
| 34 | acgttgctcataatgtctca |
| 35 | agttgtgtccatttcttcat |
| 36 | atttccgcaaaacgcgttga |
| 37 | aatcttcatcatcttcctct |
| 38 | tcagaacacaagttttggca |

| | |
|---|---|
| 39 | acttattcgcgcaaacgtga |
| 40 | gatcagattatcgactgaca |
| ***GFP* smFISH probe set** | |
| Probe # | Probe Sequence (5' to 3') |
| 1 | aagttcttctcctttactca |
| 2 | gaattgggacaactccagtg |
| 3 | cccattaacatcaccatcta |
| 4 | cctctccactgacagaaaat |
| 5 | gtaagttttccgtatgttgc |
| 6 | gtagttttccagtagtgcaa |
| 7 | acaagtgttggccatggaac |
| 8 | ggtatctcgagaagcattga |
| 9 | tcatgccgtttcatatgatc |
| 10 | gggcatggcactcttgaaaa |
| 11 | ttctttcctgtacataacct |
| 12 | gttcccgtcatctttgaaaa |
| 13 | tgacttcagcacgtgtcttg |
| 14 | taacaagggtatcaccttca |
| 15 | ataccttttaactcgattct |
| 16 | gtgtccaagaatgtttccat |
| 17 | gtgagttatagttgtattcc |
| 18 | gtctgccatgatgtatacat |
| 19 | ctttgattccattcttttgt |
| 20 | ccatcttcaatgttgtgtct |
| 21 | atggtctgctagttgaacgc |
| 22 | cgccaattggagtattttgt |
| 23 | gtctggtaaaaggacagggc |
| 24 | aagggcagattgtgtggaca |
| 25 | tcttttcgttgggatctttc |
| 26 | tcaagaaggaccatgtggtc |
| 27 | aatcccagcagctgttacaa |
| 28 | tatagttcatccatgccatg |
| ***lad-2* smFISH probe set** | |
| Probe # | Probe Sequence (5' to 3') |
| 1 | gtgcagcaaatacctgagtt |
| 2 | gggggaccaatactattttt |
| 3 | gcctgattgttctagaagct |
| 4 | ctaaatgataccacacctct |
| 5 | gtttttgaggagatcctcat |
| 6 | cctcgcaacgaagtgtcaac |
| 7 | ggagtatctgtcagtattct |
| 8 | caatgacgaatggctcaccg |
| 9 | tcccaaagaacatcttcgga |

| | |
|---|---|
| 10 | tgatccagattgactttctt |
| 11 | ccttgatgactagcatgagg |
| 12 | gttcgaaagtgctgttccaa |
| 13 | cagtgatccaagcctcaaat |
| 14 | acatcccgttttggaaagtg |
| 15 | cttccttcactctcaatagt |
| 16 | tgcaattcaacgtcagcgac |
| 17 | agccagacaattttcggatc |
| 18 | aggaatcgtctagggatcta |
| 19 | ctccggatagtttcgattac |
| 20 | accttcgttatcgacagtta |
| 21 | tctgatagctcaacggagga |
| 22 | agcacattcgtataccagag |
| 23 | tcgatattctcctctgagta |
| 24 | aatatccaactgtatccggt |
| 25 | acggttacttcactaggact |
| 26 | gccaaatatgcactgaagct |
| 27 | ctagattttggaagctctcc |
| 28 | aggatctaccgaaatcgctt |
| 29 | cgcatcatcgggatgaacat |
| 30 | ctgtatgaaccaaatgccgg |
| 31 | cagaatggagcggccataac |
| 32 | tatcacgaggtgggtcgaac |
| 33 | gtactttcttctggtaggga |
| 34 | ccagcaagacattcaagttc |
| 35 | ttatgattggagtcggctgg |
| 36 | aattttccattcatactcca |
| 37 | gaatcttctgccaattcgtg |
| 38 | ttctaccgtgatcgagtaga |
| 39 | gattcaaatttctgacgcgt |
| 40 | cattggtagactccagtatc |
| 41 | aacatatcccaacggattgc |
| 42 | aacggtgcatgagctctaac |
| 43 | tagaatggagcaccactttc |
| 44 | acgtcacaatccaggactac |
| 45 | atcaacccatcgaaccattg |
| 46 | ttatttttcccctctacaac |
| 47 | gtctgggatgtctatttgtg |
| 48 | aaatggctggctttgaggta |
| ***ser-2* smFISH probe set** | |
| Probe # | Probe Sequence (5' to 3') |
| 1 | ctgcaccgaatcagtgtaat |
| 2 | gagtcgatggctcgtaacac |

| | |
|---|---|
| 3 | cgtttataacactatcccgg |
| 4 | caacgtcgtcgtgctgactg |
| 5 | gtttcatagaggtcatggga |
| 6 | aaccagctcgacagttggaa |
| 7 | ccaagtacgtaattgttcct |
| 8 | ccgacaactgtcattgcaat |
| 9 | ctgaacactgcgacgacgac |
| 10 | tgaactttttcagcggtcg |
| 11 | atgctgcaagtgagacgaga |
| 12 | aaggcatgacgaatatcgcc |
| 13 | tttaccacccgctagaaaag |
| 14 | gtcagcagtggtgaagaatt |
| 15 | ttgagaattgatgacgtgca |
| 16 | atggcccaatatcgatctaa |
| 17 | tgcatagttgattggattgt |
| 18 | acgaatttggtcgtacgttt |
| 19 | catccaatgattggaggcac |
| 20 | tcatattctcctgccaatta |
| 21 | gtagacaacccgcaactatc |
| 22 | cggcagcagagaagacaaca |
| 23 | actaggagtgggaggaagaa |
| 24 | tttgacatacaccactacca |
| 25 | gttggcgagcactgatgaaa |
| 26 | cactgcgtcctcgattagtg |
| 27 | ggcattctggattctcatta |
| 28 | atcgacatctttcggtagtc |
| 29 | gaagtgcgtgctgattctac |
| 30 | ttcgtggtttcatcttttgg |
| 31 | gcacgatccattattgtgat |
| 32 | tcacatccttcactttcact |
| 33 | tttggatttccttcatcctc |
| 34 | ttttcgcaggactgcagtag |
| 35 | ttcgccacactgattttctc |
| 36 | agcaatagtttttgctgctc |
| 37 | atggtagccaacagaatgaa |
| 38 | atggtcgtatgacatatgct |
| 39 | gcatgaagtttacatgtttc |
| 40 | acgtgaaagcctgttcaact |
| 41 | gtgaagagttgatatagcca |
| 42 | ataccgtaaaggaatgggtt |
| 43 | ctcggcgaaactccaaattt |
| 44 | agggcacagaatcttcttga |
| 45 | tctcctctgctctaatacag |

| | |
|---|---|
| 46 | taaggttgcgcactcattcg |
| ***zig-3* smFISH probe set** | |
| Probe # | Probe Sequence (5' to 3') |
| 1 | agcagcatttttccaaatgc |
| 2 | tgcagcaaggacagatatgc |
| 3 | tctctcctgatgacaatgga |
| 4 | ccaaatttgagactgctgca |
| 5 | tgagtggagtcgatttctcg |
| 6 | tcagggatggtttagtagtt |
| 7 | tgctaactgtgttgtcttca |
| 8 | atcgaagagttacgctctcg |
| 9 | cagtcggtgtggacagtacg |
| 10 | tccttttcccagtaaatgac |
| 11 | cattaagctctttatctccc |
| 12 | cccatagcgttcaatacttt |
| 13 | gatacctgactcgactgtag |
| 14 | ggatttggtagctactggtg |
| 15 | atatgatggagatttgcgca |
| 16 | gtggctacacacttatagga |
| 17 | cagagctttcgacggtatcg |
| 18 | gtttgtccttcaactgagat |
| 19 | gcgcgttgatttacacttaa |
| 20 | attgtgattactggggctga |
| 21 | agttcaaatcgggattcagt |
| 22 | gcagataagagtagctgcat |
| 23 | aattggctctacgatcagca |
| 24 | cttgtcttcgaacatccaat |
| 25 | ggccgctgtcaaaatctatt |
| 26 | accggatggaagtagttcat |
| 27 | ctggattttccggatgagaa |
| 28 | agtatgaacccatgtctgac |
| 29 | ctccatacttgttatgagca |
| 30 | acaagaaggtttcaccacgg |
| 31 | gcaatatgttttttggtggg |
| 32 | aggggtacaattcaacagga |
| 33 | tgttcacgggtttgtaagga |
| 34 | tgcatagaatcaagtgttgt |
| ***ast-1* smFISH probe set** | |
| Probe # | Probe Sequence (5' to 3') |
| 1 | gtggctgacgagacgacttg |
| 2 | tacttgatcacatcctcatc |
| 3 | tttgatggtttgaatggccg |
| 4 | ttgtaggacggaagagcagt |

| | |
|---|---|
| 5 | gcattccattgaaaaatggg |
| 6 | cgatttggcgggaaatcgtt |
| 7 | tgtgttgtcattgtagagca |
| 8 | gatcattttcacttttctgc |
| 9 | gagaggttcataccggtgaa |
| 10 | tcctgatgatgaagccgatg |
| 11 | ctgatcctttgaagatgtgg |
| 12 | cggtatagaattgacggcgg |
| 13 | atttccagagctattactgc |
| 14 | tacttgagctcgatccattg |
| 15 | aagacgtcacttttgctttc |
| 16 | gaacgcgttcatagagatgt |
| 17 | tttttgatcctggagtggcg |
| 18 | ctcggaatgttatgatcctc |
| 19 | tgtagtaggagctgagcatg |
| 20 | gagttactcagcttgagagc |
| 21 | gtctggattggcaaacgatg |
| 22 | ttggtcccaggatttgatag |
| 23 | cggaatgtgcgaggtttttg |
| 24 | gaaattgccagagctgagtc |
| 25 | ttgtccgagaggagttcaag |
| 26 | ggtaatcacttcggagtagc |
| 27 | gatccacaagcttgaactct |
| 28 | cattttcgagccacttcatc |
| 29 | gtttggctttgactttcttt |
| 30 | agctctcgacatcttatcat |
| 31 | ggacttttgccattatattc |
| 32 | ttatacgcatatctcttccc |
| 33 | cctgtgcgattccttgaaaa |
| 34 | gtggatgagaagcagttgga |
| 35 | ccccatcgcatgagaattaa |
| 36 | acgatgagaaatcaggagca |
| 37 | gatcgataatttgcacttgt |
| 38 | attaaacccggcaatattca |
| 39 | ggttgaaaattgtgctgccg |
| 40 | ctccaaatgcactgtagttg |
| 41 | ccgagagattattcgaggtg |
| 42 | taaagagggaatgctcgtgc |
| 43 | ggaattatggggggtgacta |
| 44 | ggtaggagagatcatagatt |
| 45 | agaaatcccgttcacttcaa |
| 46 | ctcaggagactagatggaca |
| 47 | agaggtatgtgcaaaaccaa |

| | |
|---|---|
| 48 | atctagagacattcgaggag |
| ***D2089.3* smFISH probe set** | |
| Probe # | Probe Sequence (5' to 3') |
| 1 | gaggctcgatatatatgtct |
| 2 | aggactgagtgtgtctgtaa |
| 3 | agaagggcacacggaaatga |
| 4 | gttgatgatcactgaacaca |
| 5 | atatagtgtgttgcgtgttg |
| 6 | ccgcgtagaatgtagtatca |
| 7 | ctctcttgtacctgattacc |
| 8 | ttcaccgtgttttatataca |
| 9 | gatctctgatgtgtaagtgc |
| 10 | tgaagatcagcatcgtacca |
| 11 | aacagctctccgatgacaag |
| 12 | attaaccaaatgtgctcctc |
| 13 | cactgatcgaaagtccacca |
| 14 | aggtcagcgtcgatgttcaa |
| 15 | cactggagttagcttcttga |
| 16 | acgcgatgagttctgctaat |
| 17 | aattgacggtaacgttgggc |
| 18 | acacaactcgcacacgactc |
| 19 | gtcgacacattctccacaac |
| 20 | ggcgagttgcacacaaattc |
| 21 | acagagatcagtggatccag |
| 22 | tccgtgtgaattgaaaccgt |
| 23 | ctggctttctttgtttttca |
| 24 | atcgcttggcaaactgtttg |
| 25 | atcgttttctaacatctgga |
| 26 | tgtttgtaatcgaggtggga |
| 27 | cgtgacttgtttatttgaca |
| 28 | gctggggacaaacatgatct |
| 29 | agatcattggaacatgtgtt |
| 30 | taacattgactgccggtaga |
| 31 | tcattgtccaaatgctcaca |
| ***flp-12* smiFISH probe set** | |
| Probe # | Probe Sequence (5' to 3') |
| 1 | CCTCCTAAGTTTCGAGCTGGACTCAGTGgtttactgaaagttcagctt |
| 2 | CCTCCTAAGTTTCGAGCTGGACTCAGTGgggcgatgatgacttgaaca |
| 3 | CCTCCTAAGTTTCGAGCTGGACTCAGTGtgttgcaataaggcagaaca |
| 4 | CCTCCTAAGTTTCGAGCTGGACTCAGTGctttgactttctgtgtagca |
| 5 | CCTCCTAAGTTTCGAGCTGGACTCAGTGcgggagtacttcaggcgatc |
| 6 | CCTCCTAAGTTTCGAGCTGGACTCAGTGatcttgtttacgctttcgtg |
| 7 | CCTCCTAAGTTTCGAGCTGGACTCAGTGtagagcattcagcagctgag |

| | |
|---|---|
| 8 | CCTCCTAAGTTTCGAGCTGGACTCAGTGtgaagtgcttccaactctga |
| 9 | CCTCCTAAGTTTCGAGCTGGACTCAGTGcgtttttcggccatcttgag |
| 10 | CCTCCTAAGTTTCGAGCTGGACTCAGTGtgaactcgaacttgttgcgt |
| 11 | CCTCCTAAGTTTCGAGCTGGACTCAGTGtgctctactttcgtccaaat |
| 12 | CCTCCTAAGTTTCGAGCTGGACTCAGTGacaatgctcggctcaaatga |
| 13 | CCTCCTAAGTTTCGAGCTGGACTCAGTGagatccttttcaaggttgga |
| 14 | CCTCCTAAGTTTCGAGCTGGACTCAGTGgattcagagattcagaaggt |
| 15 | CCTCCTAAGTTTCGAGCTGGACTCAGTGgctatcacaattcattacga |
| 16 | CCTCCTAAGTTTCGAGCTGGACTCAGTGaagttttattttcatcggct |
| ***nlp-80* smiFISH probe set** | |
| Probe # | Probe Sequence (5' to 3') |
| 1 | CCTCCTAAGTTTCGAGCTGGACTCAGTGacgatttatgacggccattc |
| 2 | CCTCCTAAGTTTCGAGCTGGACTCAGTGgcttcggataaagcaaacag |
| 3 | CCTCCTAAGTTTCGAGCTGGACTCAGTGccaactccatgcgactgtaa |
| 4 | CCTCCTAAGTTTCGAGCTGGACTCAGTGaatcgcgacatctgcatgcg |
| 5 | CCTCCTAAGTTTCGAGCTGGACTCAGTGctccttttaccggttcttga |
| 6 | CCTCCTAAGTTTCGAGCTGGACTCAGTGgtagtggatgtacggatcac |
| 7 | CCTCCTAAGTTTCGAGCTGGACTCAGTGcatgggacgtccgaaatact |
| 8 | CCTCCTAAGTTTCGAGCTGGACTCAGTGatatgtcgagccagccgaat |
| 9 | CCTCCTAAGTTTCGAGCTGGACTCAGTGtttcgtttacaacgactccg |
| 10 | CCTCCTAAGTTTCGAGCTGGACTCAGTGagacacggggaaattatgtt |
| 11 | CCTCCTAAGTTTCGAGCTGGACTCAGTGagttgggtttttgtgaatgt |
| 12 | CCTCCTAAGTTTCGAGCTGGACTCAGTGattttcgcatcaatctcatt |
| 13 | CCTCCTAAGTTTCGAGCTGGACTCAGTGcattcaatccatgacgagca |
| 14 | CCTCCTAAGTTTCGAGCTGGACTCAGTGtatatagacctcctgcaagt |
| 15 | CCTCCTAAGTTTCGAGCTGGACTCAGTGctaggtatccggaaaagaca |
| ***C15C7.4* smiFISH probe set** | |
| Probe # | Probe Sequence (5' to 3') |
| 1 | CCTCCTAAGTTTCGAGCTGGACTCAGTGggcagaccagtacaattgaa |
| 2 | CCTCCTAAGTTTCGAGCTGGACTCAGTGatcagtttgtggagctgtag |
| 3 | CCTCCTAAGTTTCGAGCTGGACTCAGTGggaattgagaggagacgcga |
| 4 | CCTCCTAAGTTTCGAGCTGGACTCAGTGgcgctgatagtttgatcagt |
| 5 | CCTCCTAAGTTTCGAGCTGGACTCAGTGtgaacattctgtctctgtcg |
| 6 | CCTCCTAAGTTTCGAGCTGGACTCAGTGttgatactccaccatatcga |
| 7 | CCTCCTAAGTTTCGAGCTGGACTCAGTGtctctccttttcttattgg |
| 8 | CCTCCTAAGTTTCGAGCTGGACTCAGTGtcatcggcatttggtgtata |
| 9 | CCTCCTAAGTTTCGAGCTGGACTCAGTGtacaagctgctcgtcaacag |
| 10 | CCTCCTAAGTTTCGAGCTGGACTCAGTGtagaattgaccgtggcgatc |
| 11 | CCTCCTAAGTTTCGAGCTGGACTCAGTGgttgctcgcaatacatttca |
| 12 | CCTCCTAAGTTTCGAGCTGGACTCAGTGacaacgcgtttgaccgtctc |
| 13 | CCTCCTAAGTTTCGAGCTGGACTCAGTGgggatttcgaatgtttcaga |
| 14 | CCTCCTAAGTTTCGAGCTGGACTCAGTGagtaaaatggcatcctggac |
| 15 | CCTCCTAAGTTTCGAGCTGGACTCAGTGgtttcgttatacttgaagtc |

| | |
|---|---|
| 16 | CCTCCTAAGTTTCGAGCTGGACTCAGTGcttcataacatccctggata |
| 17 | CCTCCTAAGTTTCGAGCTGGACTCAGTGaagtctttggagacaccgtc |
| 18 | CCTCCTAAGTTTCGAGCTGGACTCAGTGctttgcattcatcgttgcaa |
| 19 | CCTCCTAAGTTTCGAGCTGGACTCAGTGggcgtaactcttgacttcaa |
| 20 | CCTCCTAAGTTTCGAGCTGGACTCAGTGcgactgggtatttcttgacg |
| 21 | CCTCCTAAGTTTCGAGCTGGACTCAGTGtgatcggcagcagcagaatg |
| 22 | CCTCCTAAGTTTCGAGCTGGACTCAGTGgatactgcattgcaatggga |
| 23 | CCTCCTAAGTTTCGAGCTGGACTCAGTGgcagcatcaagtagtttctt |
| 24 | CCTCCTAAGTTTCGAGCTGGACTCAGTGtgctgacgtaacgttcgtat |
| 25 | CCTCCTAAGTTTCGAGCTGGACTCAGTGatgtgtatgacgattttggt |
| 26 | CCTCCTAAGTTTCGAGCTGGACTCAGTGttgttgtgatgcttgtggtg |
| 27 | CCTCCTAAGTTTCGAGCTGGACTCAGTGagctcatcgtctctttatta |
| 28 | CCTCCTAAGTTTCGAGCTGGACTCAGTGagaagctgctgaatctatcc |
| 29 | CCTCCTAAGTTTCGAGCTGGACTCAGTGatcaatttactcggtcagtt |
| 30 | CCTCCTAAGTTTCGAGCTGGACTCAGTGtggtgggaacaattattgtt |
| 31 | CCTCCTAAGTTTCGAGCTGGACTCAGTGacggagcgagtagcatgaat |
| 32 | CCTCCTAAGTTTCGAGCTGGACTCAGTGgagttttgagtttcccgaaa |
| 33 | CCTCCTAAGTTTCGAGCTGGACTCAGTGaaatgatggctgtggcgaag |
| 34 | CCTCCTAAGTTTCGAGCTGGACTCAGTGgagaacgtgcaagttttaca |
| 35 | CCTCCTAAGTTTCGAGCTGGACTCAGTGttatggtttgtactgtgaga |
| ***nlp-20* smiFISH probe set** | |
| Probe # | Probe Sequence (5' to 3') |
| 1 | CCTCCTAAGTTTCGAGCTGGACTCAGTGacttgcatttttttggggtac |
| 2 | CCTCCTAAGTTTCGAGCTGGACTCAGTGcgatgagcagaagaagagca |
| 3 | CCTCCTAAGTTTCGAGCTGGACTCAGTGattgcgatgcagcaaaagca |
| 4 | CCTCCTAAGTTTCGAGCTGGACTCAGTGcaattcagaatcatcatccg |
| 5 | CCTCCTAAGTTTCGAGCTGGACTCAGTGaaatcgttcgttgttgctca |
| 6 | CCTCCTAAGTTTCGAGCTGGACTCAGTGcgcaattcaagatcacgagc |
| 7 | CCTCCTAAGTTTCGAGCTGGACTCAGTGgcacttctctttgcaaatgc |
| 8 | CCTCCTAAGTTTCGAGCTGGACTCAGTGatgacaacatctgcatctcc |
| 9 | CCTCCTAAGTTTCGAGCTGGACTCAGTGttcatgagcttgtggtccac |
| 10 | CCTCCTAAGTTTCGAGCTGGACTCAGTGaaagcaaagcgcataccagc |
| 11 | CCTCCTAAGTTTCGAGCTGGACTCAGTGctttggagctcgccttttg |
| 12 | CCTCCTAAGTTTCGAGCTGGACTCAGTGcgtgcaaatcgggcaaactc |
| 13 | CCTCCTAAGTTTCGAGCTGGACTCAGTGgttgatttacgcgaagctgg |
| 14 | CCTCCTAAGTTTCGAGCTGGACTCAGTGactcgtttatgacgtcatct |
| 15 | CCTCCTAAGTTTCGAGCTGGACTCAGTGcaaatttcaaatagccgccc |
